# Central extended amygdala neurons in the development of activity-based anorexia

**DOI:** 10.1101/2022.11.10.515977

**Authors:** Wesley Ilana Schnapp, JungMin Kim, Yong Wang, Sayujya Timilsena, Caohui Fang, Haijiang Cai

## Abstract

Anorexia nervosa (AN) is a serious psychiatric disease characterized by restricted eating, fear of gaining weight, as well as excessive exercise, and also is often comorbid with emotional disorders such as anxiety and depression. However, the etiology of AN is unknown and the neural mechanism that leads to AN remains to be determined. Here, we show that a specific subpopulation of neurons, marked by the expression of protein kinase C-delta (PKC-δ), in two nuclei of the central extended amygdala (EAc)— central nucleus (CeA) and oval region of bed nucleus of stria terminalis (ovBNST)—regulates development of activity-based anorexia (ABA), the current best animal model of AN. Specifically, simultaneous dual ablation of CeA^PKC-δ^ and ovBNST^PKC-δ^ neurons prevents the key phenotypes of ABA: increases in wheel activity, decreases in food intake, and life-threatening body weight loss. However, ablating PKC-δ neurons in CeA or ovBNST alone is not sufficient to prevent ABA. Correspondingly, activation of PKC-δ neurons in one type of nuclei continues to suppress food intake even when PKC-δ neurons in the other nuclei are silenced. Consistent with their role in suppressing food intake when activated, these PKC-δ neurons show increased activity with ABA development. Together, our study illuminates how neurons in the amygdala regulate ABA development and supports the complex and heterogenous etiology of AN.

## INTRODUCTION

Anorexia nervosa (AN) is a prevalent eating disorder that has the highest mortality rate of any psychiatric disorder^1,2^ and is primarily seen in females. AN is characterized by self-starvation, intense fear of gaining weight, and excessive exercise, but also is often co-diagnosed with other psychiatric and emotional disorders, such as depression, anxiety and obsessive compulsive disorder^2–8^. These characteristics of AN suggest that extensive interaction between the neural circuits regulating eating behavior and the neural circuits regulating emotion might exist to control AN development. Interestingly, AN has been associated with elevated activity in the most well established region for emotional control: the amygdala^9^. However, the specific underlying neural mechanisms involved in AN, particularly how neurons in the amygdala regulate the development of AN, remain to be determined.

Our previous work revealed that neurons expressing protein kinase C-delta (PKC-δ) in two distinct nuclei of the central extended amygdala (EAc)—the central nucleus of the amygdala (CeA) and the oval region of the bed nucleus of the stria terminalis (ovBNST)—suppress food intake when acutely activated^10,11^. Additionally, PKC-δ neurons in both regions are activated by specific anorexigenic signals: the CeA^PKC-δ^ neurons by satiety, visceral malaise nausea, and aversive taste sensation of bitter, and the ovBNST^PKC-δ^ neurons by inflammatory signals related to sickness such as interleukin-1-beta (IL-1β), lipopolysaccharides (LPS), and tumor necrosis factor (TNF-α). Independently silencing the PKC-δ in each of these regions attenuates the anorexigenic effect caused by the respective signals. These studies— combined with known amygdala hyperreactivity in human AN—support the plausibility that neural circuits involving EAc^PKC-δ^ neurons contribute to mechanisms underlying AN development. Thus, the current study investigates how this particular subpopulation of neurons in the CeA and ovBNST may regulate the development AN using the currently best animal model of AN, activity-based anorexia (ABA), in which rodents develop life-threatening self-starvation and hyperactivity tendencies when exposed to a restricted feeding schedule in combination with ad libitum access to a running wheel^12–14^.

Here we show that bilateral, simultaneous ablation of CeA^PKC-δ^ and ovBNST^PKC-δ^ neurons prevents mice from developing ABA. Specifically, elimination of these neurons inhibits the drastic reduction in body weight, insufficient food intake, and hyperactivity phenotypes, which are key characteristics in the progression of ABA development^15–18^. Ablation of PKC-δ neurons in only one of the nuclei, however, was not sufficient to prevent ABA. Consistent with these results, we found that food intake remains suppressed if the PKC-δ neurons in only one of the nuclei are silenced while the PKC-δ neurons in the other nucleus are activated. Our data also demonstrate an increased number of activated CeA^PKC-δ^ and ovBNST^PKC-δ^ neurons in mice after ABA, further confirming their role in ABA development. Additionally, we found ablation of these neurons minimized the difference in susceptibility to develop ABA between male and female mice. Together, our study demonstrates that neurons in the amygdala play a critical role in anorexia development.

## RESULTS

### CeA^PKC-δ^ and ovBNST^PKC-δ^ neurons regulate ABA development

To investigate how CeA^PKC-δ^ or ovBNST^PKC-δ^ neurons might play a role in the development of AN, we assessed how mice respond to the ABA paradigm when this subpopulation of neurons was ablated. We first focused only on female mice because human AN is observed and diagnosed disproportionally more in females^19^ and because female mice more consistently developed ABA in our mouse line and paradigm.

To ablate the PKC-δ neurons, we stereotaxically injected PKC-δ-Cre mice with a Cre-dependent adeno-associated virus (AAV) expressing caspase (AAV2-FLEX-taCasp3) bilaterally into either the CeA or ovBNST—or into both simultaneously (CeA + ovBNST) (Fig. 1a, c). We injected wild-type (WT) mice with the same virus for use as controls. Following behavior experiments, we evaluated and confirmed ablation success with immunohistochemistry and cell quantification of the labelled neurons (Fig. 1a-d). Comparison of body weight pre- and three weeks post-virus injection demonstrated that ablation itself did not significantly impact body weight (Extended Data Fig. 1a). Interestingly, even though acute silencing of CeA^PKC-δ^ and ovBNST^PKC-δ^ neurons increased food intake in previous studie^s10,11,^ chronic ablation in either or both nuclei did not increase total daily food intake, as determined before any food restriction was enforced (Extended Data Fig 1b, e). Additionally, there was no significant difference in total daily wheel activity or body mass in mice with PKC-δ neuron ablation compared to WT mice before any food restriction (Extended Data Fig. 1c-d, f-g. These baseline data indicate that EAc^PKC-δ^ neuron ablation does not induce significant changes in body weight, food intake, or wheel activity when the mice are in matching conditions with ad libitum access to wheel and food.

**Figure 1.**
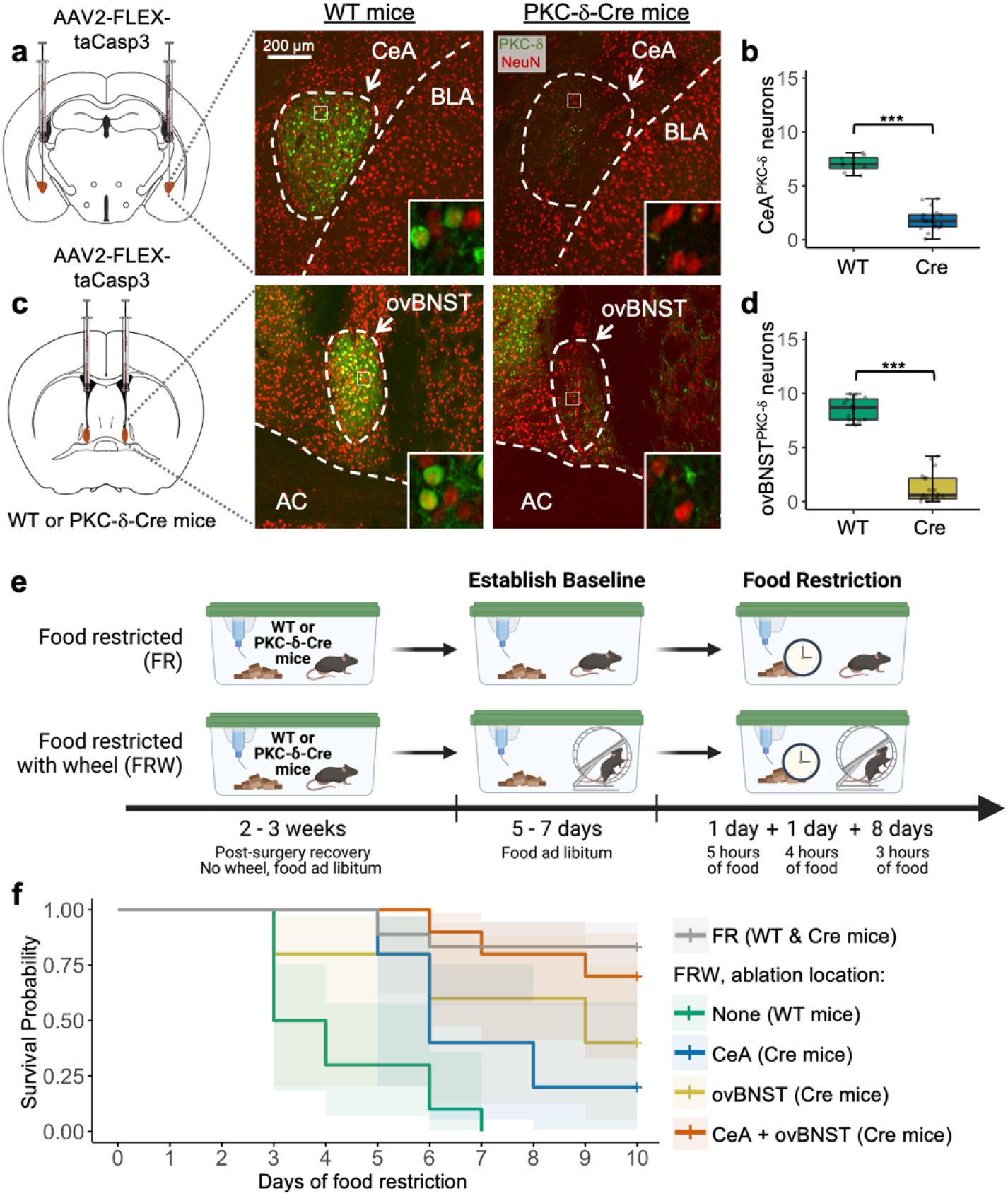
Simultaneous elimination of PKC-δ neurons in the CeA and ovBNST prevents development of ABA. **a, c,** Left: diagram illustrating bilateral injections of a Cre-dependent virus expressing caspase into the CeA **(a)** and ovBNST **(c)** regions of mouse brains. Right: representative histology of WT (PKC-δ neurons intact) and PKC-δ-Cre (PKC-δ neurons ablated). **b, d,** Quantification: average density of CeA^PKC-δ^ **(b)** and ovBNST^PKC-δ^ **(d)** neurons (number of neurons per micron x 10^4^). Unpaired t-tests; CeA-WT n = 8, CeA-Cre n = 20, ovBNST-WT n = 12, ovBNST-Cre n = 19 mice. **e,** Timeline of ABA protocol (created with BioRender.com). **f,** Survival analysis of all experimental mice. Log-rank test with respect to FR (n = 18): FRW WT/no ablation (n = 10, *p* < 0.001), FRW CeA^PKC-δ^ ablation (n = 5, *p* = 0.0084), FRW ovBNST^PKC-δ^ ablation (n = 5, *p* = 0.050), and FRW ovBNST^PKC-δ^ + CeA^PKC-δ^ ablation (n = 10, *p* = 0.50). BLA: basolateral amygdala, AC: anterior commissure, FR: food restricted (includes mice from each ablation group), FRW: food restricted with wheel. All samples are females. * *p* < 0.05, ** *p* < 0.01, *** *p* < 0.001.

After three weeks post-surgery for recovery time and gene expression, we applied the ABA assay^20^, in which food was given for a limited time each day for 10 consecutive days (Fig. 1e). Food restricted only (FR) included all mice that did not have a wheel and are a standard control group for the ABA assay because they have been shown to drop body weight but survive the limited feeding schedule^13^. In contrast but as expected, all WT mice in the food restricted with wheel group (FRW) developed ABA within the 10 days (Fig. 1f). The criteria for ABA was the point at which mice weighed 20% less than their baseline body weight (before food restriction) for two consecutive days. While ablation of PKC-δ neurons in either the CeA or ovBNST offered some level of protection from ABA for FRW mice, simultaneous ablation in both regions (CeA + ovBNST) was required for consistent prevention of ABA (Fig. 1f). FRW mice with this dual ablation showed a significant difference in survival compared to WT-FRW mice, and no significant difference compared to FR controls (Fig. 1f). These data suggest bilateral dual ablation of PKC-δ neurons in two nuclei of the EAc—the CeA and the ovBNST—have a compounding effect to regulate development of ABA.

### Dual ablation of CeA^PKC-δ^ and ovBNST^PKC-δ^ neurons prevents the reduced body weight and food intake in ABA

To determine how the mice with dual ablation of CeA^PKC-δ^ and ovBNST^PKC-δ^ neurons are able to survive the ABA conditions, we first assessed body weight change across days of the experiment, as well as corresponding food intake. Consistent with previous studies^21^, our baseline data showed that the presence of a running wheel with ad libitum food did not decrease weight loss across days (Extended Data Fig. 1d). FR mice, however, did decrease their body weight across days, but not typically to the life-threatening point requiring removal from the experiment (Fig. 1f, 2a-b). Therefore, the FR group functioned as a control in our study. As expected, there were significant differences in body weight loss between WT-FRW mice (no ablation) and their respective FR controls (Fig. 2a). However, in contrast, Cre-FRW and - FR mice (dual ablation) followed almost identical trends across all 10 days (Fig. 2b). Correspondingly, food intake was clearly disrupted and irregular in WT-FRW mice, decreasing across days as the mice lost weight (Fig. 2c), while Cre-FRW food intake very closely matched that of the respective Cre-FR control group (Fig. 2d). Individual sample data plots comparing WT-FRW (no ablation) mice to Cre-FRW mice (dual ablation) further demonstrate that mice with dual ablation were less susceptible to developing ABA (Extended Data Fig. 2a, b). Specifically, Cre-FRW body weights tended to level out at a survivable point (less than 20% loss from baseline), in contrast to the FRW-WT mice, who exhibited extreme decreases in body weight during the first few days of food restriction, to the point of requiring removal from the experiment in order to prevent death (“terminal removal”; Extended Data Fig. 2a). Additionally, Cre-FRW mice tended to gradually increase their food intake across days of limited food time exposure, and especially more consistently compared to WT-FRW mice (Extended Data Fig. 2b). In summary, these data show how FRW mice with CeA^PKC-δ^ + ovBNST^PKC-δ^ neuron ablation behaved more similarly to their respective FR controls, with adaptive increases in food intake and eventual weight stabilization, thus demonstrating increased “resilience”^22,23^ to ABA.

**Figure 2.**
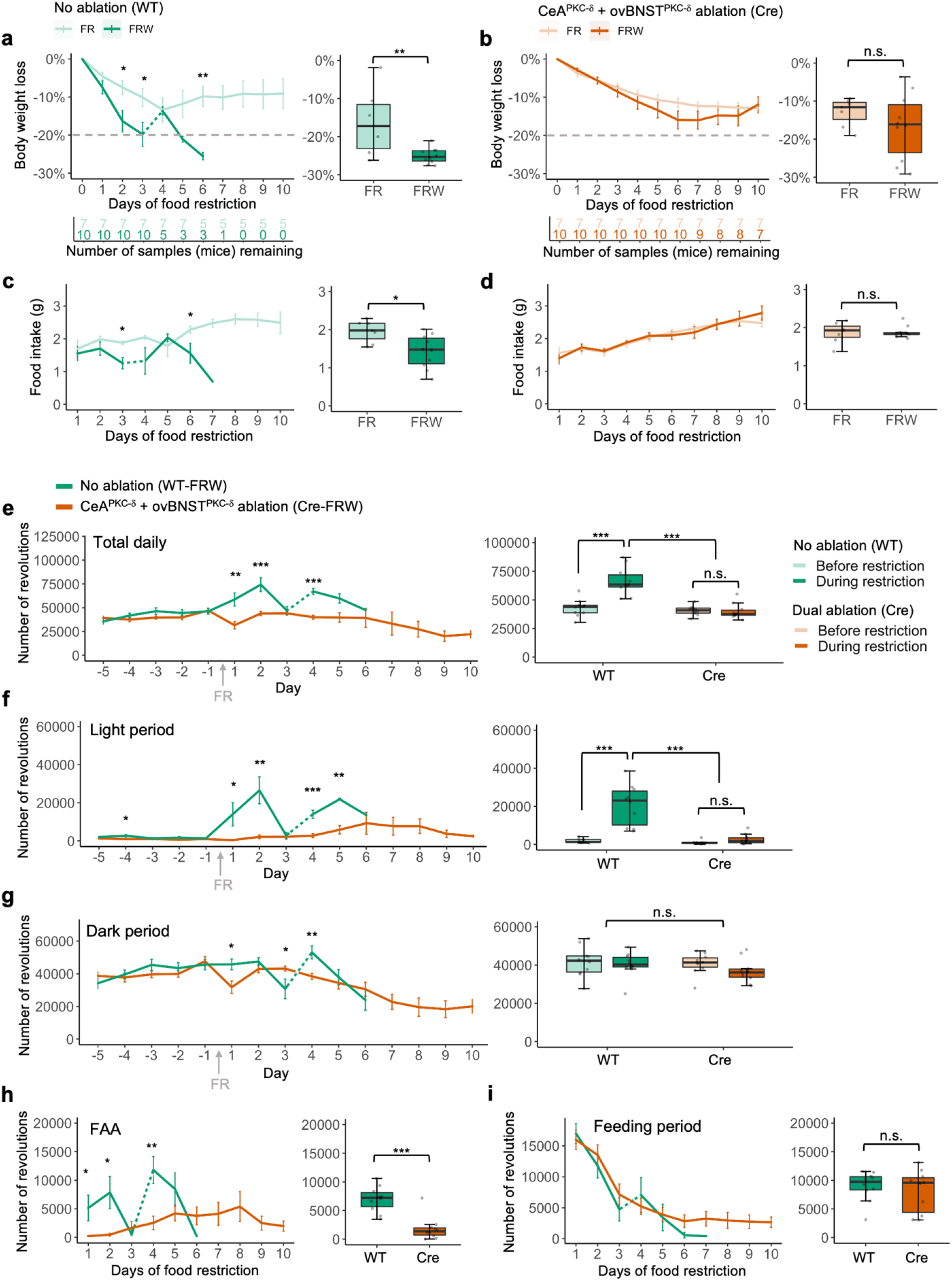
Characteristics of ABA development are mitigated with dual ablation of EAc^PKC-δ^ neurons. **a-b,** Left: population mean line plots of body weight loss across days of food restriction. Right: weight loss measurement on day of removal from experiment; either when the ABA criteria is reached (20% loss from baseline weight two days in a row) or after day 10. Gray dashed line at 20% weight loss indicates the point at which mice have developed ABA and need to be removed from the experiment to prevent death. Sudden change in the line plots for WT mice (i.e., days 3-4) is due to the removal of half the initial number of mice from the experiment in one day; indicated by dashed section in line. Unpaired *t*-tests; WT-FR n = 7, WT-FRW n = 10 **(a)**, Cre-FR n = 7, WT-FRW n = 10 **(b)**. **c-d,** Left: total food intake during each day’s feeding period. Right: average food intake across days 2-6 of the experiment (restriction began after day 1 feeding period). **e-g,** Total wheel revolutions for WT-FRW mice and Cre-FRW mice per day **(e)**, throughout light period (4am - 4pm) **(f)**, and throughout dark period (4 pm - 4 am) **(g)**. Left: population mean line plots each day during baseline (food ad libitum, days −5 to −1) and during food restriction (days 1-10). Right: average total revolutions before (days −5 to −1) and during initial food restriction (days 1-5 for total daily and light period, days 1-6 for dark period). Two-way ANOVA with Tukey HSD post-hoc analysis used for box plots comparing WT and Cre, before and during food restriction. **h-i,** Total wheel revolutions during food anticipatory activity (FAA; 4 hours preceding presentation of food) **(h)** and feeding periods **(i)**. Left: population mean line plots each day of food restriction. Right: average total revolutions during initial food restriction (days 1-5 or 6, respectively). Box plot averages based on days when more than one WT sample remained in the experiment. Arrow indicates day in which food restriction (FR) was enforced. All samples are females. * *p* < 0.05, ** *p* < 0.01, *** *p* < 0.001.

### Dual ablation of CeA^PKC-δ^ and ovBNST^PKC-δ^ neurons prevents the light period hyperactivity in ABA

Another crucial element to the development of ABA is increased wheel activity (i.e., hyperactivity)^24,25^. Daily running wheel activity in WT (no ablation) mice was not significantly different compared to Cre mice (dual ablation) *before* food restriction began (Fig. 2e). However, once food was restricted (indicated by the arrow in Fig. 2e), WT-FRW mice significantly increased their daily wheel activity, while Cre-FRW mice did not have significant alterations in daily wheel activity (Fig. 2e, Extended Data Fig. 2c-e). WT-FRW mice demonstrated hyperactivity across days of food restriction until terminal removal, whereas Cre-FRW mice tended to show relatively steady wheel activity, suggesting adaptation to food restriction parameters (Extended Data Fig. 2c).

To investigate details of the wheel activity, we examined different time frames for each day: light period, dark period, food anticipatory activity period (FAA, defined as four hours preceding food presentation^26,27^), and the limited feeding period. WT-FRW mice demonstrated extreme wheel hyperactivity during the light period (“rest/sleep” time), including FAA, yet this hyperactivity was absent in Cre-FRW mice (Fig. 2f, h, see Extended Data Fig. 3a-b for individual sample plots). Additionally, Cre-FRW mice demonstrated a trending decrease of wheel activity during the dark period (“awake” time) across days of food restriction (day 2 vs. day 10, *p* = 0.00019, unpaired *t*-test), suggesting a level of arousal adaptation that corresponds with the new feeding schedule (Fig. 2g, Extended Data Fig. 3c; statistic not shown in figure). Consistent with previous results^25^, time series data show that WT-FRW mice had strong wheel activity disruptions and abnormalities—with hyperactivity even during the light period (“rest/sleep” time)—one or two days before terminal removal (Extended Data Fig. 3e, see arrows for examples). In contrast, Cre-FRW mice showed consistent patterns of day/night wheel activity, with moderate activity during the dark period and minimal activity during the light period (Extended Data Fig. 3f). We did not see a significant difference in the wheel activity during feeding time (Fig. 2i, Extended Data Fig. 3d).

Together, these data demonstrate that Cre-FRW mice with dual ablation of PKC-δ neurons CeA and ovBNST survived ABA conditions primarily by preventing wheel activity increase upon the introduction of the new feeding schedule, as well as by gradually increasing food intake across days.

### ABA causes more EAc^PKC-δ^ neurons to be activated in response to food

Our results thus far indicate that PKC-δ neurons in both the CeA and ovBNST contribute to the development of ABA. Previous studies demonstrated that activation of CeA^PKC-δ^ or ovBNST^PKC-δ^ neurons suppresses food intake^10,11^. Correspondingly, we hypothesized that FRW mice have increased activity of CeA^PKC-δ^ and ovBNST^PKC-δ^ neurons after ABA development. To investigate the involvement of the PKC-δ neuron activity in the four different brain regions (bilateral CeA and bilateral ovBNST) in mice developing ABA, we monitored c-Fos expression in WT-FRW mice and their respective controls, WT-FR only mice, in response to food intake during the ABA paradigm. Interestingly, there were a few FRW mice in this cohort that did not develop ABA within the 10-day experiment, but they were still collected for analysis purposes (FRW non-ABA). On the day in which FRW mice reached >20% body weight loss from baseline (ABA), they were perfused 90 minutes after presentation of food. FR (Non-ABA) mice were similarly collected on comparable days to the FRW mice. The FRW (Non-ABA) mice were collected on day 10, the last day of the experiment. Double immunostaining for c-Fos^+^ and PKC-δ^+^ neurons revealed that FRW (ABA) mice had significant increases in both CeA^PKC-δ^ and ovBNST^PKC-δ^ neurons expressing Fos compared to FR (non-ABA) and compared to FRW (non-ABA) mice (Fig. 3a-b, e-f). This increase was observed in both the right and left hemispheres of the CeA and ovBNST, suggesting a bilateral importance (Extended Data Fig. 4a-b). The total number of Fos-expressing neurons in these nuclei, however, was not significantly different (Fig. 3c, g). Additionally, simple linear regression analysis indicates a significant negative correlation between food intake and c-Fos^+^ PKC-δ^+^ neurons across groups (Fig. 3d, h), which aligns with the significantly reduced food intake in FRW (ABA) mice compared to FR (non-ABA) and FRW (non-ABA) mice (Fig. 3i) on day of terminal removal. ABA resistance and susceptibility is clear when comparing the body weight loss and wheel revolutions during days of food restriction for mice in the FRW condition (Fig. 3j-m). The results here are consistent with the previously discovered function of EAc^PKC-δ^ neurons in suppressing food intake when activated^10,11^ and further support the involvement of these neurons in regulating the development of ABA.

**Figure 3.**
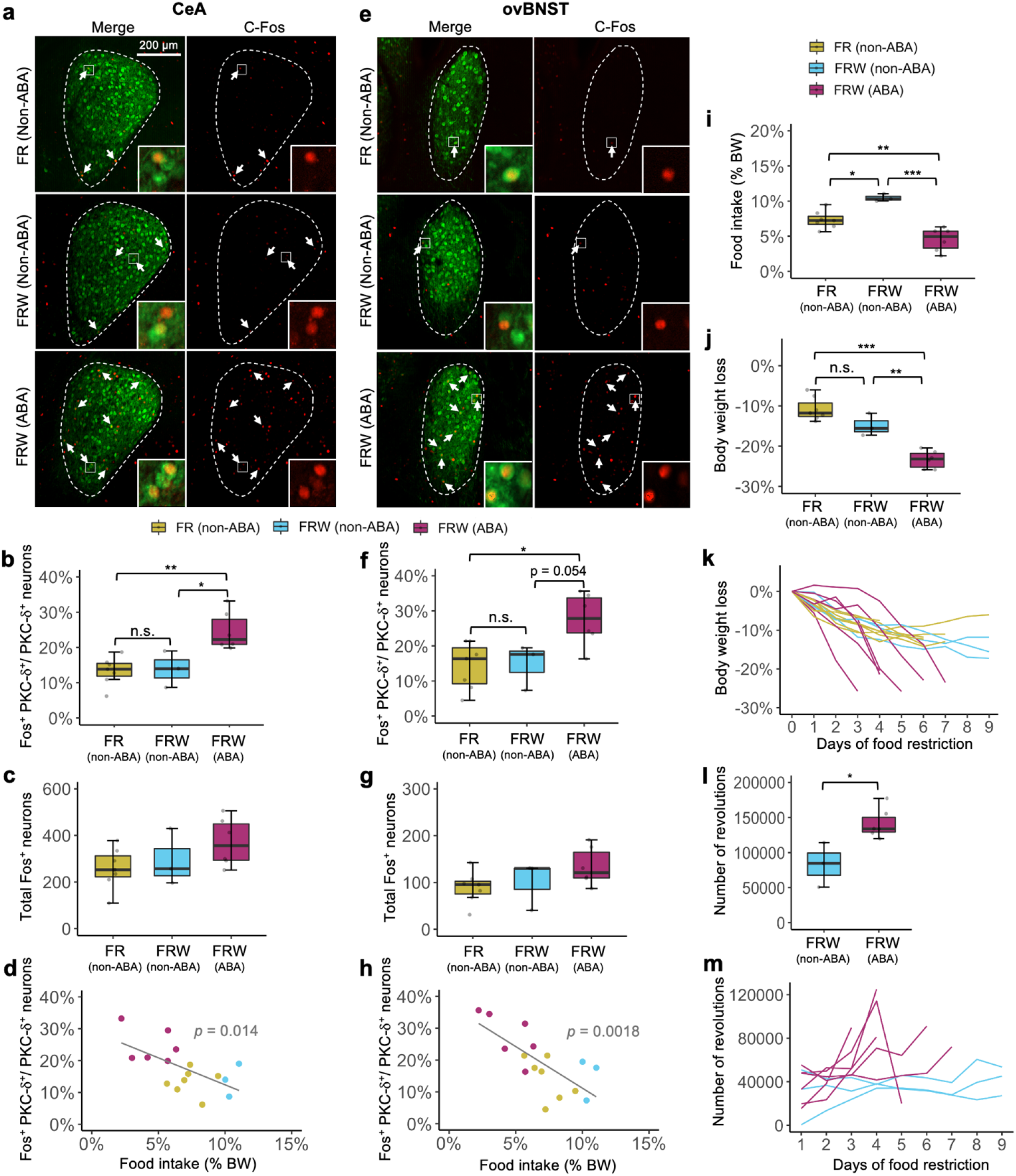
FRW mice that develop ABA show increased activity of PKC-δ neurons in both the CeA and ovBNST. **a-g,** Representative histology and quantification of Fos-like immunoreactivity in the CeA **(a-c)** and ovBNST **(e-g)** brain regions of WT mice in the food restricted only condition (FR Non-ABA; n = 7), ABA resistant mice with food restriction and wheel (FRW Non-ABA; n = 3), and ABA susceptible mice with food restriction and wheel (FRW ABA; n = 6); One-way ANOVA with Tukey HSD post-hoc. **d, h,** Relationship between Fos expression level in CeA^PKC-δ^ neurons **(d)** and ovBNST^PKC-δ^ neurons **(h)** and food intake across groups (BW = body weight); determined with simple linear regression analysis. **i,** Comparison of food intake between groups on day of removal from experiment. **j-k,** Body weight loss on day of removal **(j)** (Unpaired *t*-test) and across days of food restriction **(k)** for ABA susceptible and ABA resistant mice. **l-m,** Total wheel activity on the two full days preceding removal **(l)** (Unpaired *t*-test) and daily wheel activity across days of food restriction **(m)** for ABA susceptible and ABA resistant mice. Arrows pointing to examples of C-Fos and PKC-δ co-expression. All samples are females. * *p* < 0.05, ** *p* < 0.01, *** *p* < 0.001.

### CeA^PKC-δ^ and ovBNST^PKC-δ^ neurons function in combination

To determine how CeA^PKC-δ^ and ovBNST^PKC-δ^ neurons might exert their function at the circuit level, we examined downstream projections from the specific PKC-δ neurons in each of these discrete regions simultaneously using Cre-dependent viral tracing techniques (Fig. 4a). We alternated between EYFP and mCherry to account for innate differences in fluorescence, and found that fluorescence was largely at the same locations and with similar intensity for both the CeA and ovBNST (Fig. 4b). CeA^PKC-δ^ neurons displayed their strongest projections, in order, at the medial part of central amygdala (CeM), extended BNST region, and ventrolateral BNST (vlBNST) (Fig. 4b). ovBNST ^PKC-δ^ neuron projections were similar, but with vlBNST being the strongest, followed by extended BNST and CeM. Both CeA^PKC-δ^ and ovBNST ^PKC-δ^ neurons showed minor terminal fluorescence at the parasubthamalic nucleus (PSTh), ventrolateral medial reticular formation (vlmRt), and lateral parabrachial nucleus (LPB) (Fig. 4b). Overall, these data demonstrate how GABAergic CeA^PKC-δ^ and ovBNST ^PKC-δ^ neurons make similar contributions to intra-circuit inhibitory connections within the EAc (Fig. 4c), as well as long-range projections to multiple brain regions, suggesting that removal of one would have some disinhibitory effect, but not as great as if both were removed.

**Figure 4.**
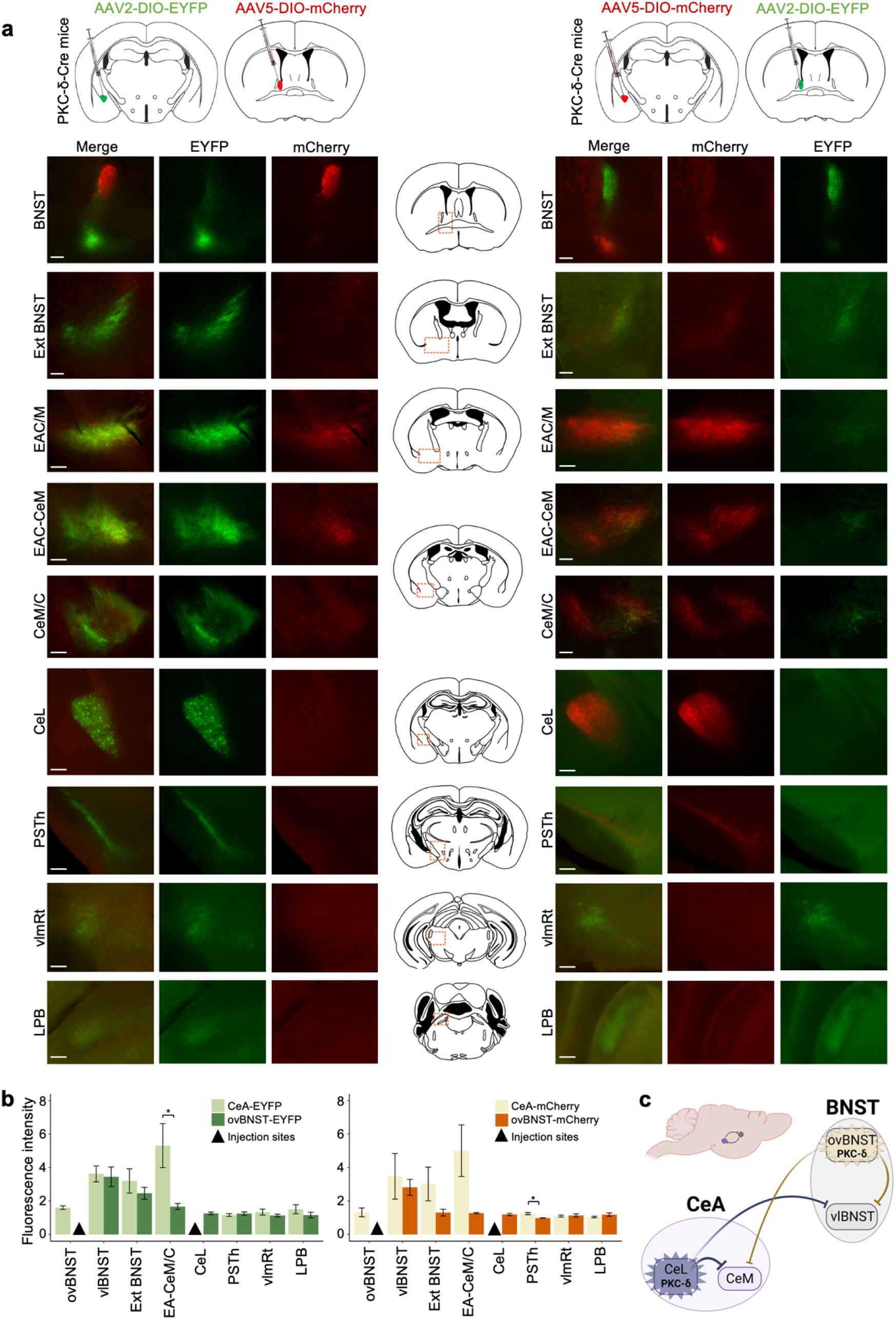
CeA^PKC-δ^ neurons and ovBNST^PKC-δ^ neurons have similar downstream targets. **a-b,** Viral tracers were used to simultaneously label PKC-δ neurons in the CeA and ovBNST, alternating with EYFP and mCherry for both. Locations and representative histology of the viral tracing **(a)**. Left: CeA^PKC-δ^ neurons tagged with EYFP and ovBNST^PKC-δ^ neurons with mCherry (n = 5). Right: CeA^PKC-δ^ neurons tagged with mCherry and ovBNST^PKC-δ^ neurons with EYFP (n = 5). 200 μm scale bars. Quantification of fluorescence intensity for each region where there are fluorescent axons and terminals **(b)**; statistical analysis with unpaired *t*-tests. Fluorescence measure not included for injection sites due to increased fluorescence intensity of cell bodies (not a proper comparison with terminals). Green and red fluorescence are graphed separately to account for innate differences in fluorescence, as EYFP is typically stronger than mCherry. **c,** Summary diagram of the intra-circuit and long-range projections from CeA and BNST PKC-δ^+^ neurons (made with BioRender.com). vlBNST: ventrolateral BNST, Ext BNST: extended BNST, EA: extended amygdala, CeM: medial central amygdaloid nucleus, CeC: capsular central amygdaloid nucleus, CeL: lateral central amygdaloid nucleus, PSTh: parasubthalamic nucleus, vlmRt: ventrolateral medial reticular formation, LPB: lateral parabrachial nucleus. * *p* < 0.05, ** *p* < 0.01, *** *p* < 0.001.

Our previous results showed that optogenetic activation of the CeA^PKC-δ^ neurons or ovBNST^PKC-δ^ neurons suppresses food intake, while chemogenetic silencing of these neurons can block the anorexia induced by the corresponding anorexigenic signals that these neurons mediat^e10,11.^ Thus, we tested whether bilateral silencing of the PKC-δ neurons in one of the two nuclei would prevent the feeding suppression caused by activation of the other nuclei. We stereotaxically injected AAV-DIO-ChR2-EYFP in one of the nuclei, as well as AAV-DIO-hM4Di-mCherry in the other nuclei; both bilaterally (Fig. 5a, c). Consistent with our previous stud^y10,11,^ chemogenetic inhibition of ovBNST^PKC-δ^ neurons increased food intake compared to saline (Fig. 5b), while inhibiting CeA^PKC-δ^ neurons had a non-significant trend of increased food intake (Fig. 5d). When PKC-δ neurons in one of the nuclei were optogenetically activated, the simultaneous silencing of PKC-δ neurons in the other nuclei caused a trend of increase in food intake (right side of graphs, saline vs. CNO), yet still significantly lower than no activation control (CNO + activation vs CNO + non-activation). These results suggest that activation of either CeA^PKC-δ^ neurons or ovBNST^PKC-δ^ neurons alone is sufficient to suppress food intake, even in the absence of activity in the other. This observation is consistent with our results that single ablation only had a mild attenuation on ABA while dual ablation prevented ABA.

**Figure 5.**
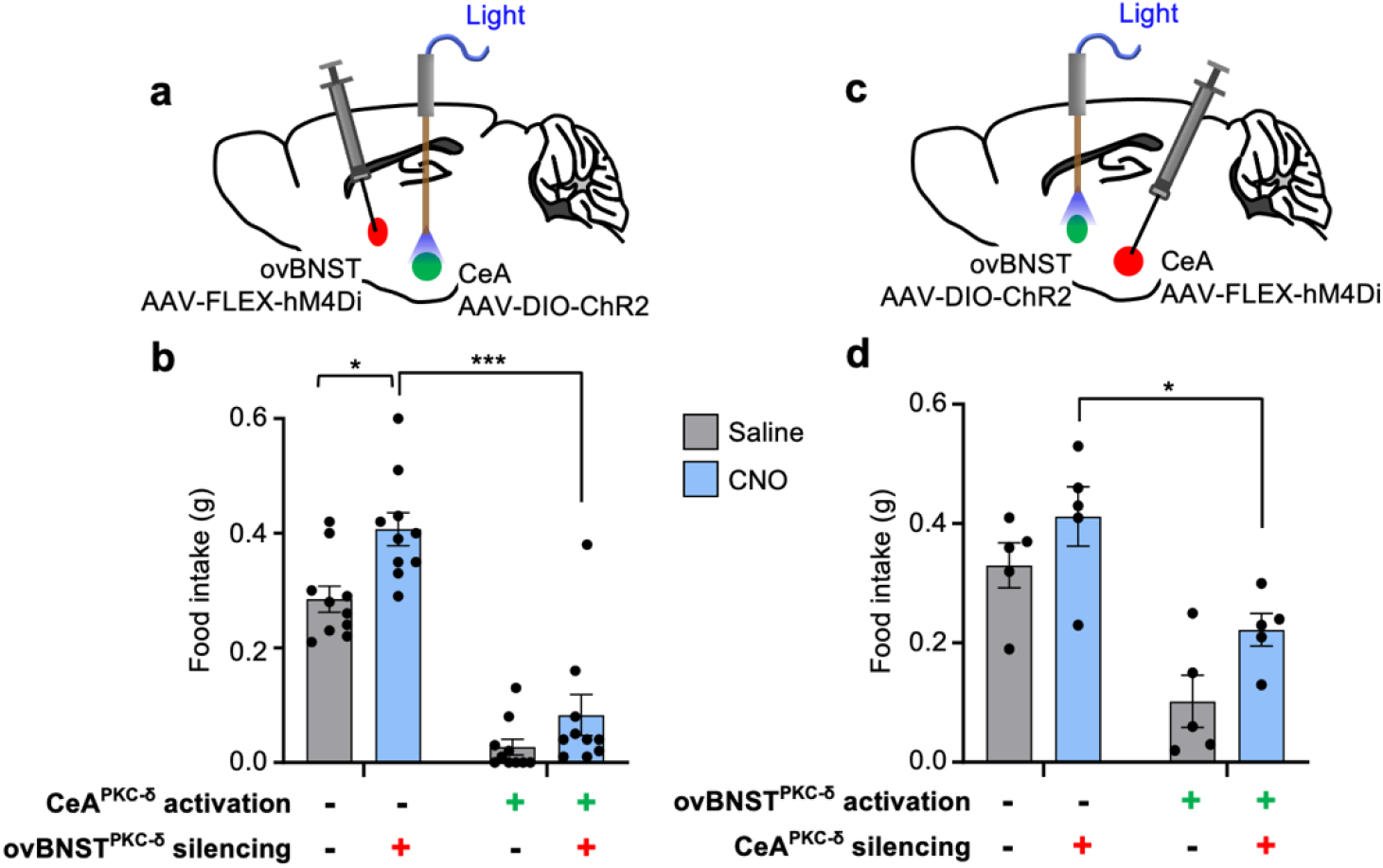
Differential food intake upon alternating simultaneous manipulations of CeA^PKC-δ^ and ovBNST^PKC-δ^ neurons. **a-d,** Diagrams outlining manipulation techniques and bar graphs displaying food intake measurements when combining optogenetic activation of CeA^PKC-δ^ neurons with chemogenetic silencing of ovBNST^PKC-δ^ neurons **(a-b)** and, vice-versa, activation of ovBNST^PKC-δ^ neurons with silencing of CeA^PKC-δ^ neurons **(c-d)**. In each bar graph, from left to right: control (saline), chemogenetic silencing only, optogenetic activation only, simultaneous optogenetic activation and chemogenetic silencing. Two-way ANOVA with Tukey’s multiple comparisons test; n = 5-10 mice in each group. * *p* < 0.05, ** *p* < 0.01, *** *p* < 0.001.

### Dual ablation of CeA^PKC-δ^ and ovBNST^PKC-δ^ neurons minimizes sexual divergence seen in ABA

Previous literature showed that behavior and survival in the ABA paradigm can vary depending on sex^16,28^. Our WT-FRW mice did indeed demonstrate sexual divergence: the females consistently developed ABA quicker and lost weight more drastically than males (Fig. 6a, c, Extended Data Fig. 5a). However, in contrast to these WT-FRW mice without ablation, Cre-FRW mice with dual ablation did not demonstrate sexual divergence in the same way. Survival analysis indicates no significant difference in probability of developing ABA, while the trend of decrease in body weight was similar between the sexes (Fig. 6b, d, Extended Data Fig. 5b).

**Figure 6.**
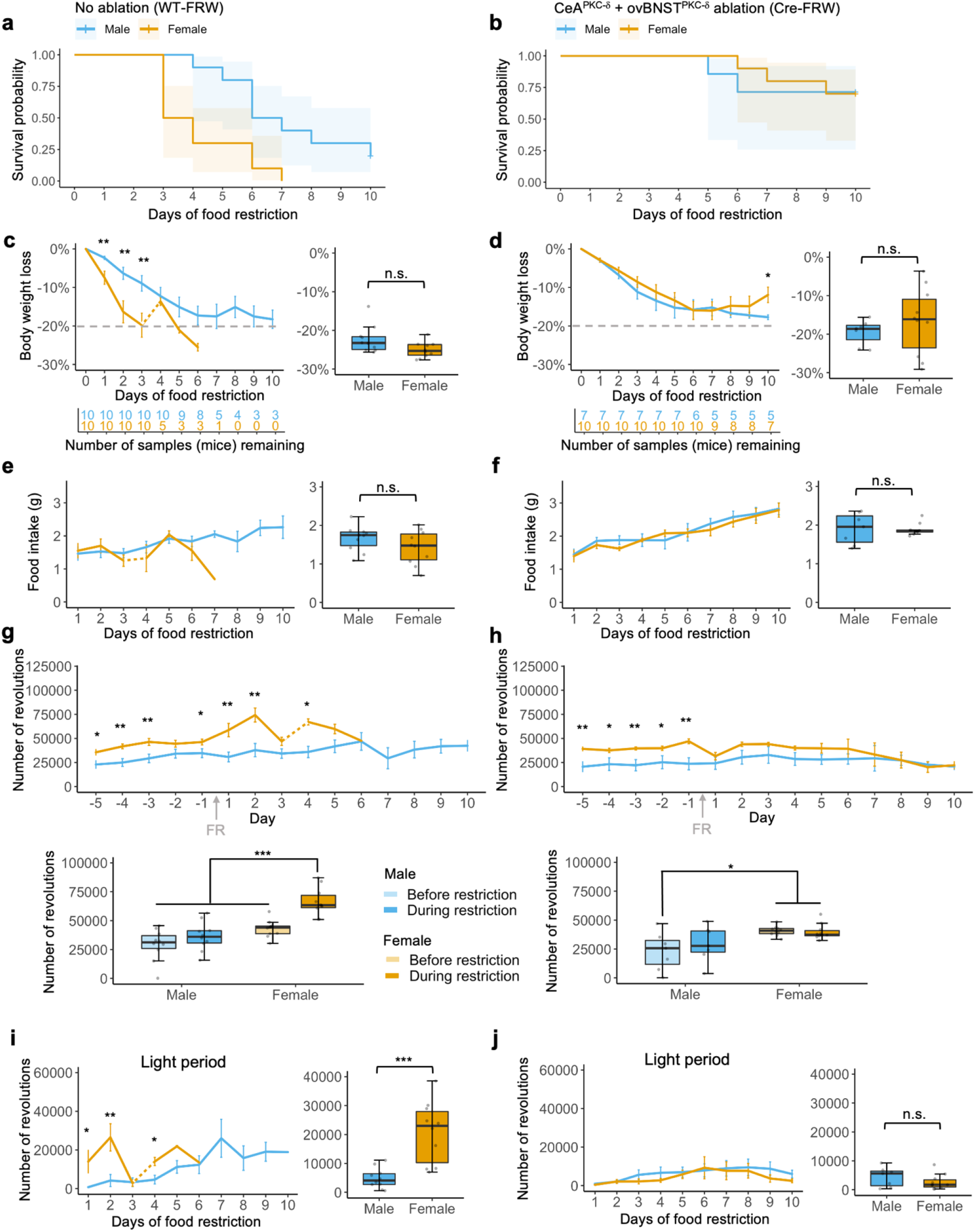
Sexual divergence in ABA is attenuated in samples with dual ablation of EAc^PKC-δ^ neurons. **a-b,** Survival analysis of WT-FRW mice **(a)** and of Cre-FRW mice **(b)** (WT *p* = 0.0045, male n = 10, female n = 10; Cre *p* = 0.93, male n = 7 and female n = 10; log-rank test). **c-d,** Left: population mean line plots of body weight loss across days of food restriction. Right: weight loss measurement on day of removal from experiment; either when the ABA criteria is reached (20% loss from baseline weight two days in a row) or after day 10. Gray dashed line at 20% weight loss indicates the point at which mice have developed ABA and need to be removed from the experiment to prevent death. Sudden change in the line plots for female WT mice (i.e., days 3-4) is due to the removal of half the initial number of mice from the experiment in one day; indicated by dashed section in line. Unpaired *t*-tests. **e-f,** Left: total food intake during each day’s feeding period. Right: average food intake across days 2-6 of experiment. **g-h,** Top: total wheel revolutions per day during baseline before food restriction (food ad libitum, days −5 to −1) and during food restriction (days 1-10). Bottom: average total wheel activity before and during initial food restriction (days 1-5). Two-way ANOVA with Tukey HSD post-hoc analysis used for box plots comparing male and female, before and during food restriction. **i-j,** Total wheel revolutions during light period (4 am - 4 pm). Left: population mean line plots across days of food restriction. Right: average total revolutions during initial food restriction (days 1-5). Box plot averages based on days when more than one WT sample remained in the experiment. Arrow indicates day in which food restriction (FR) was enforced. * *p* < 0.05, ** *p* < 0.01, *** *p* < 0.001.

While average food intake did not significantly differ between sexes in each group (Fig. 6e-f, Extended Data Fig. 5c-d), there was variance in the wheel activity (Fig. 6g-h, Extended Data Fig. 5e-f). Consistent with previous studies^29,30^, there was a slight increase in daily wheel activity in females compared to males for both groups before food restriction started (Fig. 6g-h). However, while the male and female WT-FRW mice increased their differences in wheel activity upon food restriction, the difference was minimized upon food restriction for Cre-FRW mice. Accordingly, the wheel activity during the light period was significantly different between male and female WT-FRW mice, but not with Cre-FRW mice (Fig 6i-j, Extended Data Fig. 5f-g). FAA, specifically, was also significantly different between sexes for WT-FRW mice, but not for Cre-FRW mice (Extended Data Fig. 6a-b). While dark period wheel activity was increased in females of both experimental groups compared to the respective males for the first 5 days of food restriction, the activity for Cre-FRW females decreased and plateaued to become aligned with males for the latter half of food restriction days (Extended Data Fig. 6c-d). Additionally, there was no difference in the wheel activity for the sexes of either group during the feeding period (Extended Data Fig. 6e-f). Overall, decreased wheel activity during the light period indicates decreased susceptibility to develop ABA, which is seen in a portion of the male WT-FRW mice, as well as in a majority of both male and female Cre-FRW mice with dual ablation. Therefore, these data suggest that EAc^PKC-δ^ neurons might be involved in arousal and circadian rhythm changes associated with ABA that are typically different between sexes.

## DISCUSSION

AN is a psychiatric condition that also involves disruptions in eating behavior and energy homeostasis, making it an inherently complicated disorder. Consequently, the etiology of AN remains ambiguous.

Although functional alterations of brain regions associated with AN have been observed in human neuroimaging studies^31–33^, details of neural mechanisms that might cause AN are still being uncovered. Studies exploring these underlying neural processes often use the ABA animal model, which demonstrates three major characteristics leading to the development of anorexia: life-threatening decrease in body weight, reduced food intake, and increased physical activity^13,14,18,33^. The increased running wheel activity, in particular, has been established as a robust predictor for the susceptibility of ABA^25^. In this study, we found that PKC-δ neurons in two nuclei of the central extended amygdala are necessary for mice to develop each of these key phenotypes that contribute to ABA.

Much of the previous literature studying the neural circuits underlying ABA, however, emphasizes reward systems as the primary neural dysfunction responsible for ABA development^9,34^. For example, subcutaneous infusion of a low dose of non-selective dopaminergic antagonist attenuates ABA while a high dose exacerbates it^27^. Similarly, administration of D2/3 antagonists ameliorates the ABA phenotypes in mice^35^. These results are consistent with a recent study demonstrating that a D2 to D1 shift in dopaminergic pathway regulates ABA, and that interruption or inhibition of the dopamine signaling pathways attenuates ABA^36^. However, these manipulations of the dopamine pathways tend to have only a mild attenuation effect on ABA development, with majority of the animals still developing ABA, but in a delayed manner. Surprisingly, a small—although not significant—*increase* in FAA is observed after infusion of dopaminergic antagonist^27^, which is, in fact, contradictory to being less susceptible to developing ABA. Involvement of the dopamine pathway in ABA development is also supported by an experiment showing that overexpression of dopamine receptor in nucleus accumbens (NAc) induces more weight loss, reduces food intake, and increases wheel activity during ABA^37^. Chemogenetic excitation of the rewarding pathway from ventral tegmental area (VTA) to NAc pathway in female rats attenuates weight loss and increases food intake, yet has no effect on overall running activity; rather, it induces a surprising increase in FAA^38^. Furthermore, using a mild ABA development rat model, in which ~40% of the animals develop ABA, a study showed chemogenetic silencing of the pathway from medial prefrontal cortex to NAc prevents ABA development^39^. However, this manipulation has little to no effect on food intake, does not impact overall running wheel activity and, again, causes a surprising *increase* in FAA. These studies suggest that the reward pathway may play a role in regulating ABA, but does not directly contribute to ABA development as the effect on preventing ABA is mild with some characteristics in ABA not being affected, or even being affected in the opposite direction.

Another promising neural circuit that might contribute to ABA development is in the hypothalamus region, which has been well-studied and established as regulating feeding behavior, metabolism, and energy balance^40–42^. For example, activation of neurons expressing agouti-related protein (AgRP) in the arcuate nucleus promotes eating behaviors. However, a recent study found that activation of the AgRP neurons made no difference in rescuing the decreased body weight or caloric intake, but allowed mice to sustain the increased wheel activity before physical exhaustion in the regular ABA paradigm^43^, suggesting alteration of the AgRP neuron activity does not contribute to ABA development. It remains to be determined if other neural pathways in this canonical eating center regulate ABA development.

It has long been known that AN is comorbid with emotional conditions and, therefore, that development of the disorder may be attributed to the neural circuits that control emotions, especially those in the amygdala regions^44,45^. Consistent with this theory, neuroimaging studies in humans have suggested that the function of amygdala or amygdala associated mesolimbic brain regions are altered in those with AN^31–33^. However, whether neurons in amygdala regulate ABA was not known. In fact, only recently have neurons in the amygdala, especially the CeA and BNST regions, been demonstrated to regulate eating behavior and eating suppression in anorexigenic conditions^10,11,46–51^. Among these neurons, CeA^PKC-δ^ neurons are preferentially activated by anorexigenic signals such as satiety, visceral malaise and nausea, and bitter taste, but not by LPS-induced sickness^10^. On the other hand, the ovBNST^PKC-δ^ neurons are preferentially activated by anorexia signals related to inflammation or sickness, such as IL-1β, LPS, and TNF-α, but not by the CCK satiation signal^11^. Chemogenetic silencing of these PKC-δ neurons blocks the anorexigenic effect induced by the distinct corresponding signals. Here, we demonstrated that single ablation of CeA^PKC-δ^ neurons or ovBNST^PKC-δ^ neurons had only a mild effect in attenuating ABA, while dual ablation of the PKC-δ neurons simultaneously in these two nuclei prevented ABA development. Notably, all key phenotypes were attenuated—life-threatening body weight loss, insufficient food intake, overall running wheel hyperactivity, and increased FAA—to a level indistinguishable from their respective controls (FR only or access to wheel with food ad libitum). These results clearly demonstrate that neurons in the amygdala play a critical role in ABA development. Since these PKC-δ neurons regulate emotional behaviors or have wide interactions with the neurons that regulate emotions^52,53^, future research regarding how emotions such as depression and anxiety contribute to AN development, or are affected by AN, is warranted.

While acute manipulations of the CeA^PKC-δ^ or ovBNST^PKC-δ^ neurons have been shown to affect food intake^10,11^, we did not observe significant changes in food intake or body weight after ablation of these neurons during the baseline period with food ad libitum (Extended Data Fig. 1), suggesting a possible compensation mechanism after ablation. Nevertheless, our results showed that the activity of these PKC-δ neurons increased significantly in response to food after ABA development (Fig. 3), which is consistent with their anorexigenic effect when activated, as well as the fact that ablation of these neuron can maintain sufficient food intake and body weight in the ABA paradigm. Importantly, increased wheel activity during the light period, including FAA, that is characteristic to development of ABA is prevented with dual ablation of the PKC-δ neurons. This hyperactivity behavioral response signifies disrupted circadian rhythm^21,26^, thus implying involvement of these neurons in circadian clock mechanisms and, more specifically, food-entrained oscillators^54^. This finding aligns with specific expression of circadian clock protein, PER2, in the CeA and ovBNST^55^, and changes in expression that occur with the estrous cycle^56^, thus, potentially accounting for the sex differences we observed. Future work will involve clarifying how these PKC-δ neurons and PER2 or other clock proteins may interact to regulate circadian disruption that occurs with the ABA paradigm.

Data from our neural tracing experiment showed that PKC-δ neurons in the CeA and ovBNST have similar downstream targets, which aligns with previous work indicating similar anatomical connections of the CeA and BNST neurons^57^. In combination with this evidence, the requirement of simultaneously ablating PKC-δ neurons in two brain regions to sufficiently block all key characteristics observed in ABA supports the notion of neurons in each of these distinct nuclei as having different functions, yet in a way that is integrated and coordinated. This necessity for dual ablation also suggests that ABA development involves contribution from a combination of multiple anorexigenic factors rather than a single factor. Additionally, the notion that phasic fear is associated with the CeA, while sustained fear and anxiety are identified with the BNST^58,59^—two types of stress responses that could be independently relevant in ABA and AN—further supports the significance of both regions and the heterogenous aspect of such disorders. Given that these neurons are characterized as being GABAergic^60^, ablation would lead to elimination of inhibitory projections, thus, disinhibition of downstream targets. Removing only one of the sources of inhibition may not be sufficient for complete disinhibition. Another possibility for why single ablation does not have the same effect is that the neurons in each nucleus may innervate different subpopulations, even if they project to similar brain regions, but future experiments are needed for clarification. Consistent with this concept of both nuclei being necessary, we demonstrated that food intake is still suppressed when the PKC-δ neurons in one type of nuclei are silenced if PKC-δ neurons in the other are activated (Fig. 5). Additional work should go into identifying the downstream disinhibited neurons in order to gain deeper knowledge and insight into the circuit.

In summary, our study provides evidence that malfunction of neural circuits in the amygdala—the emotion center of the brain—contributes to ABA development, and demonstrates that amygdala circuits may be a more relevant and robust therapeutic target in the treatment of AN. We also propose a multi-origin possibility for ABA development, which suggests that treating AN requires consideration of combining multiple factors or targeting multiple brain regions.

## METHODS

### Mice

We crossed PKC-δ-Cre C57BL/6 mice (generated by David Anderson’s lab^53^) with wildtype (WT) C57BL/6 mice from the Charles River Laboratory to get PKC-δ-Cre and WT littermate mice; the same mouse line as was used in previous studies from our lab^10,11^. The genotype of offspring that we generated and used from these mice were determined by PCR of genomic tail DNA. Stereotaxic survival surgery was performed when mice were 2-3 months old. All mice were housed on a 12-hours light (4 am)/dark (4 pm) cycle, with ad libitum access to water and rodent chow, except for during the ABA experiment food restriction and food intake tests. All animal care and experimental procedures were in accordance with ethical regulations, conducted according to the National Institutes of Health guidelines for animal research, and approved by the Institutional Animal Care and Use Committee (IACUC) at the University of Arizona.

### Virus and tracer

For Cre-dependent ablation, we used rAAV2-FLEX-taCasp3-Tevp, a virus generated by Dr. Nirao Shah’s lab^61^. For Cre-dependent anterograde tracing, we used rAAV2-EF1a-DIO-EYFP and rAAV5-hSyn-DIO-mCherry, generated by Dr. Karl Deisseroth’s lab and by Dr. Bryan Roth’s lab, respectively. For optogenetic activation, we used rAAV2-EF1a-DIO-ChR2-EYFP generated by Dr. Karl Deisseroth’s lab. For chemogenetic silencing, we used rAAV5-hSyn-DIO-hM4Di-mCherry generated by Dr. Bryan Roth’s lab These viral constructs were deposited and packaged into viral vectors either at the University of North Carolina (UNC) Viral Vector Core or Addgene at a titer of 4-6×10^12^ genome copies per ml. Upon arrival to our lab, the virus stocks were aliquoted and stored at −80°C until used.

### Stereotaxic survival surgery

All mouse surgeries were performed using aseptic techniques with a stereotaxic frame (Model 1900 Stereotaxic Alignment System, Kopf Instruments), as previously described^11^. Injection coordinates (in mm) relative to midline, bregma, and skull surface at bregma were followed as (x, y, z): ovBNST (±1.13, +0.30, −4.10) and CeA (±2.85, −1.40, −4.70). Viruses were microinfused through a pulled-glass micropipette with 20–50 μm tip outer diameter connected with a Nanoliter Injector (Nanoliter 2010, World Precision Instruments) at a rate of 10 units per minute (1 unit = 4 nl). After each injection, the micropipette was left in place for 5 min to allow for diffusion of the liquid, followed by withdrawal of 5 units at the same rate, 0.3 mm in the +z direction to remove any unwanted spread of virus. Injection volume for the caspase virus was 55 units (220 nl) and for the anterograde tracer was 40 units (160 nl) per injection location. The caspase virus was injected bilaterally and the virus for tracing experiments was injected unilaterally. For the optogenetic tests, optical ferrule fibers (200 μm in diameter) were implanted bilaterally ~0.5 mm above the injection coordinates. After ferrule fiber implantation, dental cement (C&B Metabond) was used to secure the fiber to the skull. For postoperative care, mice were given water with 1.2% Septra antibiotics for 5-7 days, as well as injected intraperitoneally with ketoprofen (5 mg/kg) daily for 3 days. Mice were allowed three weeks after virus injection surgery for recovery and viral expression before used for behavioral experiments or euthanized for tracing analysis.

### Activity-Based Anorexia (ABA)

Stereotaxic surgery was performed on mice undergoing the ABA protocol (based on Welch, 2018^20^) for the ablation experiment. After 2-3 weeks for recovery and gene expression, mice were individually housed, and a wheel was added to their respective home cages. Control mice without a wheel were individually housed at the same time. Food and water were given ad libitum for the next 5-7 days, while establishing a baseline of wheel activity, body weight, and food eaten. Total food and body weight of each experimental mouse was measured daily during the hour before start of light off/active period. Mice were habituated to the experimenter handling during this time, as well. MED Associates Inc low-profile Wireless Running Wheels (ENV-047) with the respective Wheel Manager Software was used for continually recording running data. After establishing the baseline and acclimating the mice to the new environmental condition for 5-7 days, we started food restriction. On the first day, mice were given ad libitum food for 5 hours, starting at the beginning of the dark cycle. On the second day, mice were given ad libitum food for 4 hours, still starting at the onset of the dark cycle period. From the third day until the end of the study, mice were given ad libitum food for 3 hours, again starting at the beginning of the dark cycle.

Each day, mice—and their respective wheels—were transferred from their home cage with bedding to an empty cage, for the sake of measuring the food consumed most accurately. More than enough pre-weighed regular chow (NIH-31, Zeigler Bros, Inc) was provided in the empty cage (~10 g total). Water was given ad libitum, both during the feeding period and during the food restriction period. After the feeding period, mice were weighed and transferred back into their original home cages with the wheel if they had one. Red light was used in the dark room to prevent disruption to the light cycle and circadian rhythm. In another room, total food left was then measured for each sample, and calculated from what was given to determine amount eaten. The empty cages were cleaned and used again for the rest of the experiment.

For ethical purposes, mice were removed from the experiment when body weight loss exceeded 20% of their respective baseline weight two days in a row, when measured before given food. If the mouse’s body weight remained under 20% after the feeding period on the first day of that measurement, it was also removed. Mice were monitored on the day after reaching 20%, and were removed from the experiment if they appeared to be in critical condition or reached 20% loss before group measurement time, in order to prevent unexpected deaths. Therefore, since mice were removed at various times on their second day of being below 20% weight loss, the final day of running wheel activity data was not included in analysis for all mice who developed ABA. Removal from the experiment (i.e., “terminal removal”) meant mice were given ad libitum food immediately, while the wheel was taken out and a cardboard house was placed in the home cage.

### Immunohistochemistry and histology

Immunostaining and histology analyses were performed in order to check (1) the level of ablation of PKC-δ neurons after the ABA experiments, (2) c-Fos expression in PKC-δ neurons, and (3) virus expression for the tracing experiments. All mice were deeply anesthetized with ketamine/xylazine and perfused with 20 ml PBS followed by 20 ml of 4% paraformaldehyde (PFA) in PBS. Mice in the c-Fos experiment were perfused 90 minutes after presentation of food. The brains were then extracted, post-fixed in 4% PFA overnight at 4°C, rinsed with PBS, and then sectioned with a vibratome (Leica, VT1000S). Brains from mice that underwent the ABA experiment were sliced at 100 μm thickness, while brains from the viral tracing were sliced at 150 μm thickness. The brain slices with the viral tracing were mounted on glass slides and imaged. The brain slices from ABA mice were stained with antibodies to tag all neurons (NeuN) and neurons expressing PKC-δ. Brain tissue slices were stained with primary antibody at 4 °C overnight, in a blocking solution containing 5% donkey serum and 0.5% Triton X-100. After three rounds of 5-10 minute washes in PBS with 0.1% Triton X-100 solution, the tissues were incubated in secondary antibodies in the PBS-0.1% Triton X-100 at room temperature for 1-2 hours. Tissue slices were then washed for three times for 5-10 minutes in PBS before being mounted on glass slides. Vectashield mounting medium was added before placing coverslips on top. Imaging was done using a ZEISS AxioZoom v16 Fluorescent Microscope with Apotome 2 Structured Illumination Module for optical sectioning.

Primary antibodies used were rabbit anti-PKC-δ (Abcam, ab182126, 1:1000), guinea pig anti-NeuN (Fisher/Sigma, ABN90MI, 1:1000), and guinea pig c-Fos antibody (SYSY, 266 308, 1:5000). Secondary antibodies used were Alexa Fluor 488 donkey anti-rabbit IgG (Jackson Immuno Research Inc. 711-545-152, 1:500) and Alexa Fluor 594 donkey anti-guinea pig IgG (Jackson Immuno Research Inc. 50-194-3535, 1:500).

### Food intake with chemogenetic silencing and optogenetic activation

After three days of habituation for at least 20 min each day, mice were food-deprived, with water provided ad libitum, one day before test. Mice were briefly anaesthetized with isoflurane and coupled with optic fibers and Clozapine-N-oxide (CNO) IP injection (Enzo life science-Biomol, BML-NS105-0005, freshly dissolved in 0.9% NaCl saline to a concentration of 1 mg/ml) at 5 mg/kg. Saline was injected as vehicle control. After at least 25 min of recovery, optogenetic activation was performed as previously described^10,11^. Food intake was then measured in a 20 min feeding session. The light was delivered by a blue laser (Shanghai DreamLaser: 473 nm, 50 mW) just after the mice were introduced into the testing cage. To be consistent with previous experiments^10,11^, 15 Hz, 10 ms light pulses were used to activate ovBNST^PKC-δ^ neurons, while 5 Hz, 10 ms light pulses were used to activate CeA^PKC-δ^ neurons. No difference in food intake were observed between male and female mice after these manipulations^10,11^, thus both male and female mice were pulled together in this experiment.

### Quantification and statistical analysis

All WT and PKC-δ-Cre mice that were injected with the taCasp3 virus (see virus and tracer section for details) were perfused, and brains were extracted for analysis of ovBNST^PKC-δ^ and CeA^PKC-δ^ cell population (see immunohistochemistry and histology sections for details). Data from WT mice that underwent food restricted (FR) and food restricted with wheel (FRW) conditions for the C-Fos experiment were collected and analyzed in the same way. For the ovBNST, 3-4 brains sections that included anterior, middle, and posterior ovBNST regions were analyzed and averaged per animal. Similarly, for the CeA, 6-8 brain sections that included anterior, middle, and posterior CeA were analyzed and averaged per animal. The number of PKC-δ cells per area of interest (ovBNST and CeA) were quantified using the multi-point tool/cell counter plug-in in

FIJI/ImageJ. Samples with partial ablation were included, as this ablation technique is not always absolutely complete, or to the same degree in the exact same parts of the region targeted (ovBNST and/or CeA). To determine and calculate the percentage of PKC-δ cells expressing C-Fos in the ovBNST and CeA, the cells simultaneously expressing PKC-δ and c-Fos were counted with the channels tool, divided by the total PKC-δ cells in the same region, and multiplied by 100.

As much of the brains as possible from the PKC-δ-Cre mice injected with tracing virus were sliced with the vibratome and imaged with the AxioZoom (see previous sections for details). The FIJI/ImageJ software was used to measure the level of fluorescence. For each region, the measurements were averaged across all slices in which they appeared. *The Mouse Brain In Stereotaxic Coordinates* ^62^ was used for reference to identify location of fluorescence. The regions were grouped and designated as follows: bed nucleus of the stria terminalis (BNST, oval and ventral-lateral; plates 29-31), extended BNST (Ext BNST, plates 32-36), extended amygdala and medial/capsular central amygdaloid nucleus (EA-CeC/M; plates 37-40), central amygdaloid nucleus (CeA, CeM, CeL, CeC; plates 41-46), parasubthalamic nucleus (PSTh; plates 46-51), ventrolateral medial reticular formation (vlmRt, plates 55-62), lateral parabrachial nucleus (LPB, plates 76-79). No difference in fluorescence was observed between male and female mice, thus sexes were combined in this analysis.

Data were analyzed with RStudio or GraphPad Prism Software. Line plot error bars represent mean ± s.e.m. Box plots show interquartile range (IQR), ranging from 25^th^ to 75^th^ percentile, with the bar in the box representing 50^th^ percentile (median). Top/bottom whiskers are the largest/smallest value within 1.5 times IQR above/below the 75^th^ and 25^th^ percentile, respectively. Outside values beyond either end of the box and error bars are greater than 1.5 times the IQR. Unpaired *t*-test was used to compare two groups with one variable. One-way ANOVA was used to compare three or more groups with one variable, and two-way ANOVA for groups with more than one variable; both with Tukey HSD post-hoc analysis. Survival analysis plots display the Kaplan-Meier estimate of time-to-event (i.e., development of ABA) with right censoring method to account for subjects that had not developed ABA by the end time point (day 10). Log-rank test was used to determine if there was a statistically significant difference in survival curves between groups. Shading around curves represent the 95% confidence intervals for the point estimates. A *p*-value less than 0.05 was considered significant.

## AUTHOR CONTRIBUTIONS

W.I.S. designed the experiments, collected, analyzed, and interpreted the data, as well as wrote the manuscript. H.C. conceived the project and helped with experiment design, data interpretation, and manuscript writing. J.K. performed pilot ABA experiments. S.T. helped with the ABA, immunohistochemistry, imaging, and analysis for the c-Fos experiment. Y.W. performed the experiment and analysis for Figure 5. C.F. managed the mouse colony, including genotyping and characterizing the mouse lines.

## ACKNOWLEDGEMENTS

We would like to thank W. Haubensak and D. Anderson for the PKC-δ-Cre mice; Ananya Nigam for helping with some of the post-ABA histology imaging and cell quantification; Jeannette Hoit, Jennifer Teske, Shivani Mann, Marco Contreras Abarca, Matthew Schmit, and Masa Miscevic for critical reading and comments on the manuscript. Additional thanks to the University of Arizona Animal Care. This research was supported by grants from the Klarman Family Foundation Eating Disorders Research Grants Program (Grant ID 4770) and the NIDDK (R01 DK124501) to H.C.

---

**Extended Data Fig. 1.**
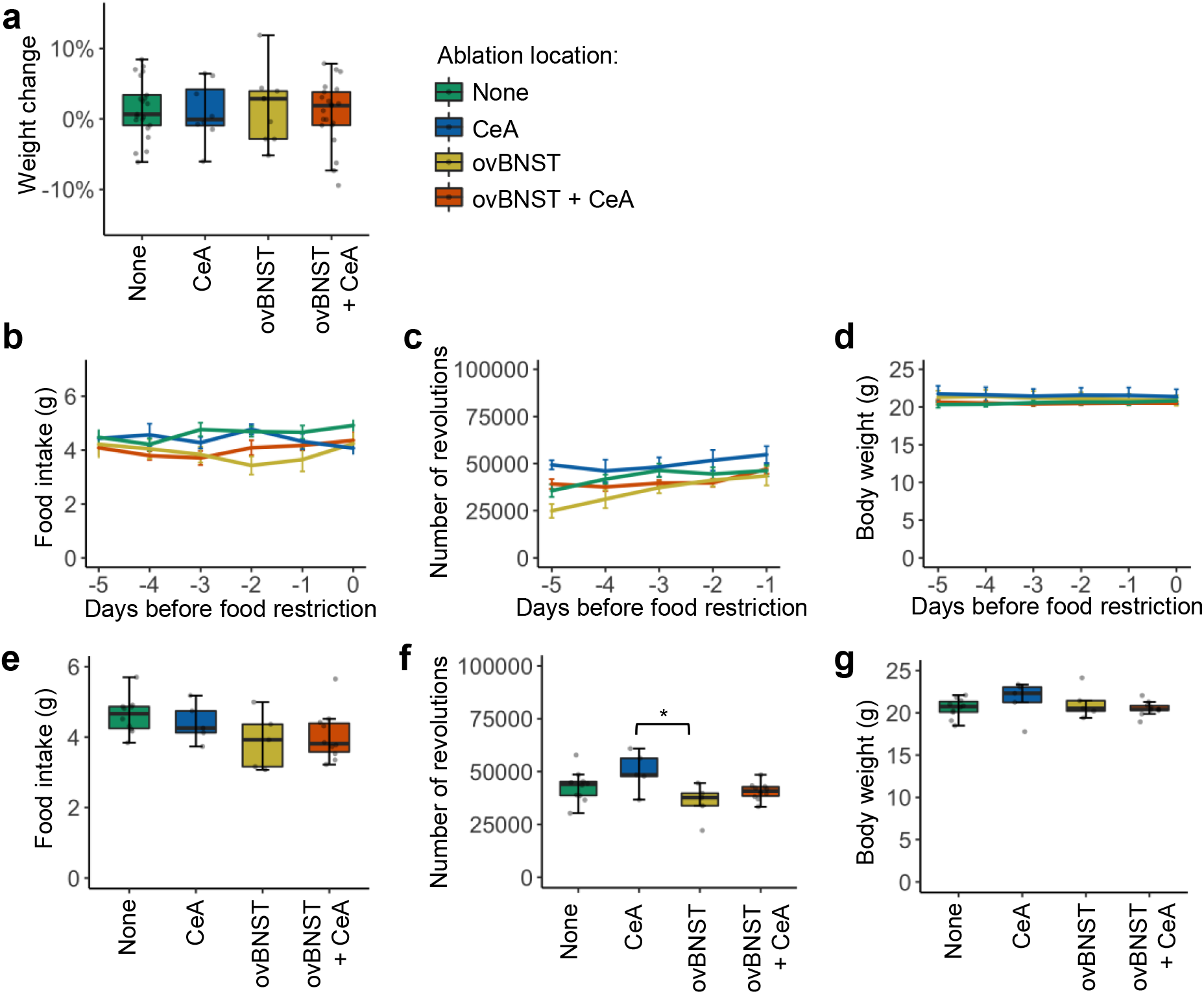
Baseline data of energy balance before ABA conditions are implemented. **a,** Body weight change two to three weeks after virus injection surgery; enough time for recovery and gene expression (None: n = 21, CeA: n = 9, ovBNST: n = 9, ovBNST + CeA: n = 21). One-way ANOVA with post-hoc Tukey HSD (*p* = 0.96). **b-d,** Population mean for each group, on each day with wheel and food ad libitum for food intake (**b**) (One-way ANOVA with post-hoc Tukey HSD: day −3, None vs. ovBNST + CeA *p* = 0.019; day −2, None vs. ovBNST *p* = 0.016, CeA vs. ovBNST *p* = 0.028), total daily number of wheel revolutions (**c**) (day −5, None vs. CeA *p* = 0.037, CeA vs. ovBNST *p* < 0.001, ovBNST vs. ovBNST + CeA*p* = 0.028), and body weight (**d**) (None: n = 10, CeA: n = 5, ovBNST: n = 5, ovBNST + CeA: n = 10). **e-g,** Average food intake (**e**), total daily number of wheel revolutions (**f**), and body weight (**g**) across five days with wheel and food ad libitum. All samples are females. * *p* < 0.05, ** *p* < 0.01, *** *p* < 0.001.

**Extended Data Fig. 2.**
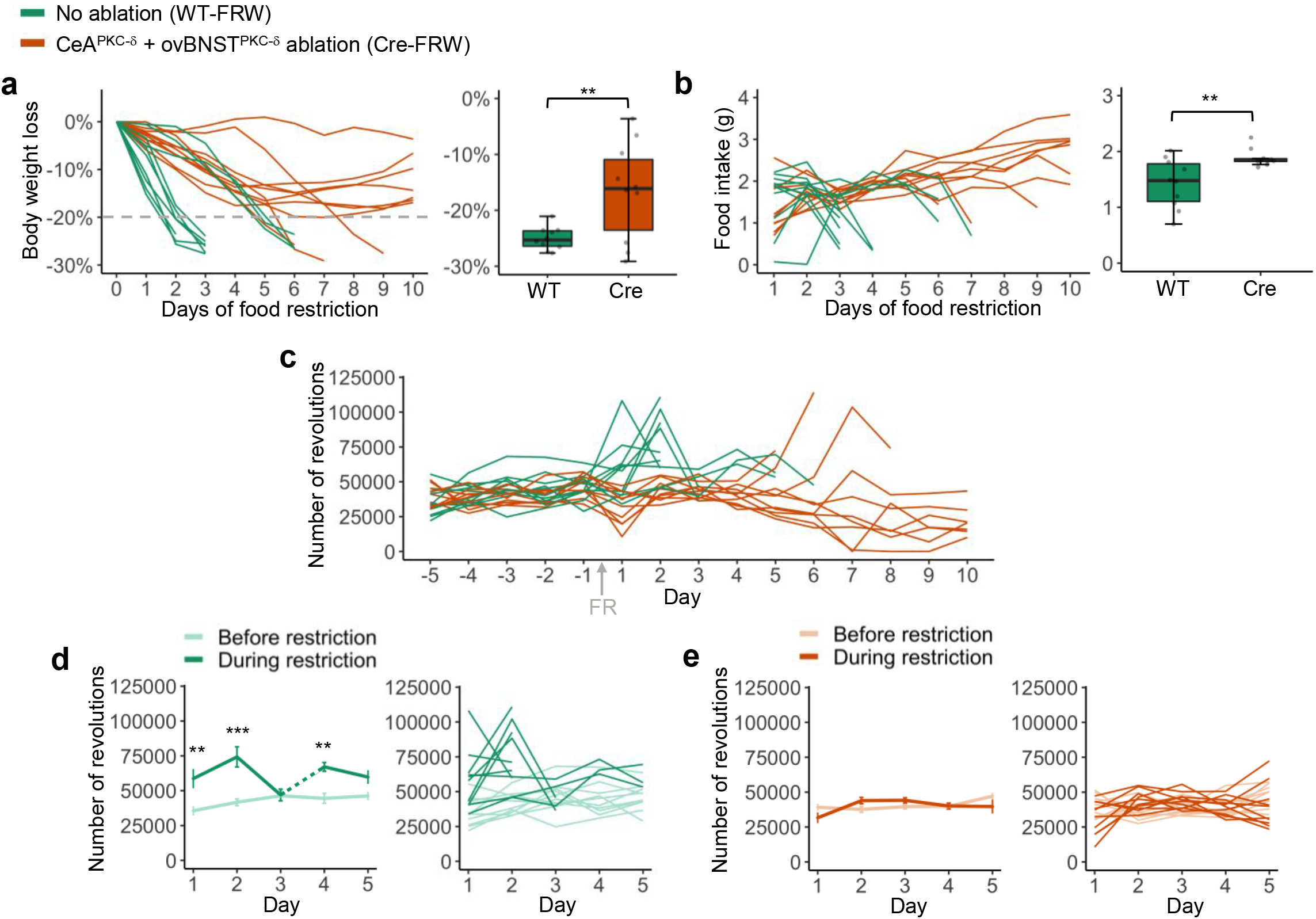
Cre-FRW mice with dual EAc^PKC-δ^ neuron ablation survive ABA by adapting food intake across days and due to lack of wheel hyperactivity. **a-b,** Body weight loss (**a**) and food intake (**b**) in ABA paradigm (FRW). Left: Line plots for individual samples. Right: weight loss measurement on day of removal from experiment (**a**) and average food intake across days 2-6 of experiment (**b**). Unpaired */*-tests. **c,** Total wheel revolutions per day during baseline (food ad libitum, days −5 to −1) and during food restriction (days 1-10). See Figure 2 for statistical differences in average total wheel activity during baseline and initial food restriction (days 1-5). **d-e,** Comparison of total daily wheel activity with food ad libitum (before) to initial days of food restriction (during) for WT (no ablation) (**d**) and Cre (dual ablation) (**e**) mice; Unpaired *t*-tests. Arrow indicates day in which food restriction was enforced. WT n = 10, Cre n = 10. \**p* < 0.05, ** *p* < 0.01, *** *p* < 0.001.

**Extended Data Fig. 3.**
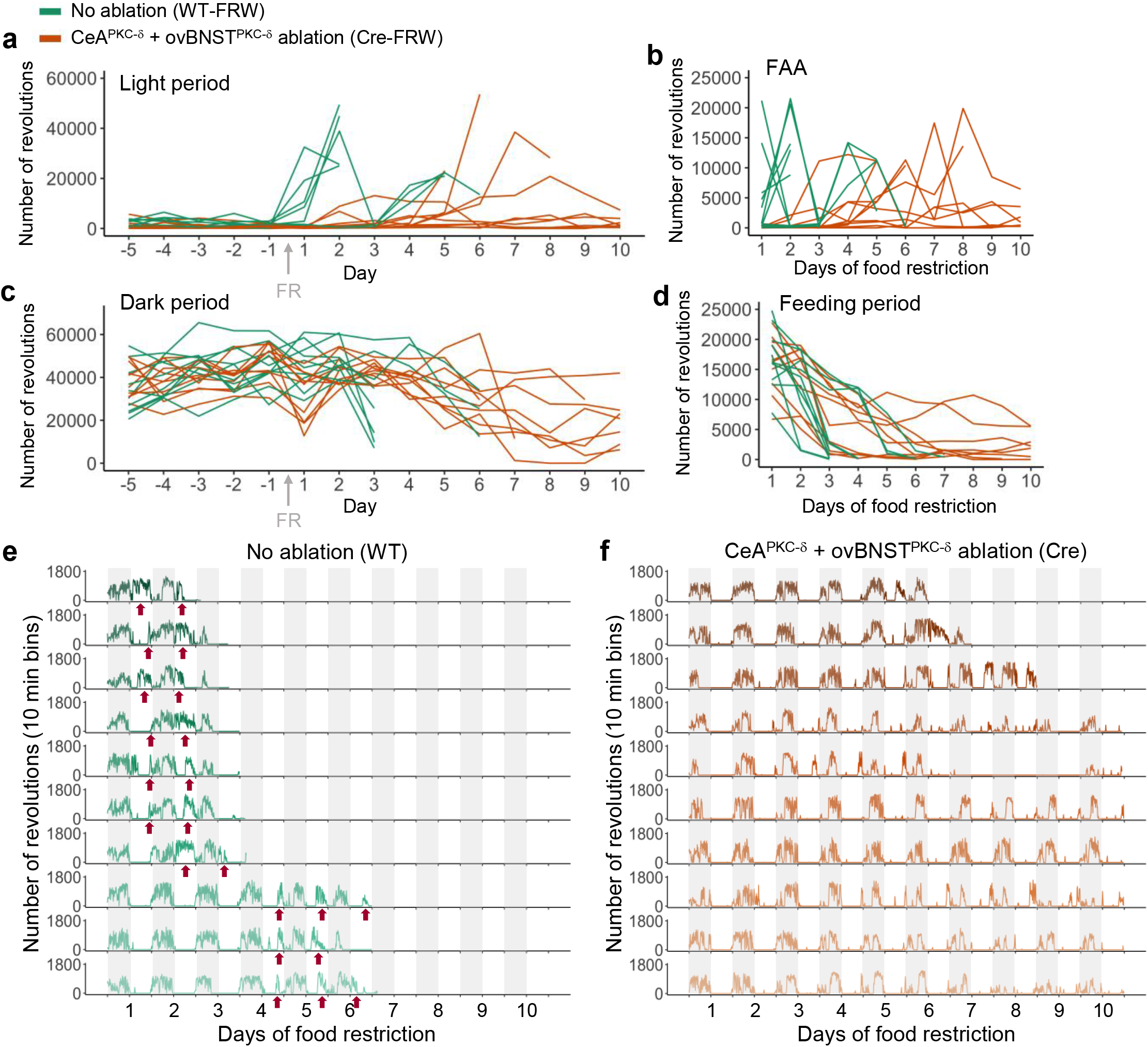
Wheel activity for majority of Cre-FRW mice with EAc^PKC-δ^ neuron ablation is not disrupted upon food restriction. **a-d,** Individual sample wheel activity displayed for total number of revolutions each day during light period (4 am – 4 pm) (**a**), food anticipatory activity (FAA, 4 hours preceding food) (**b**), dark period (4 pm - 4 am) (**c**), and feeding period (**d**). Arrows on line plot in (**a**, **c**) indicates day in which food restriction (FR) was enforced. **e-f,** Time series, split into 10 minute bins, of wheel activity across days of food restriction for each mouse. Arrows on time series plots point to ABA characteristic disrupted wheel activity for WT-FRW mice (**e**). Shaded region represents dark period, white represents light period. WT n = 10, Cre n = 10.

**Extended Data Fig. 4.**
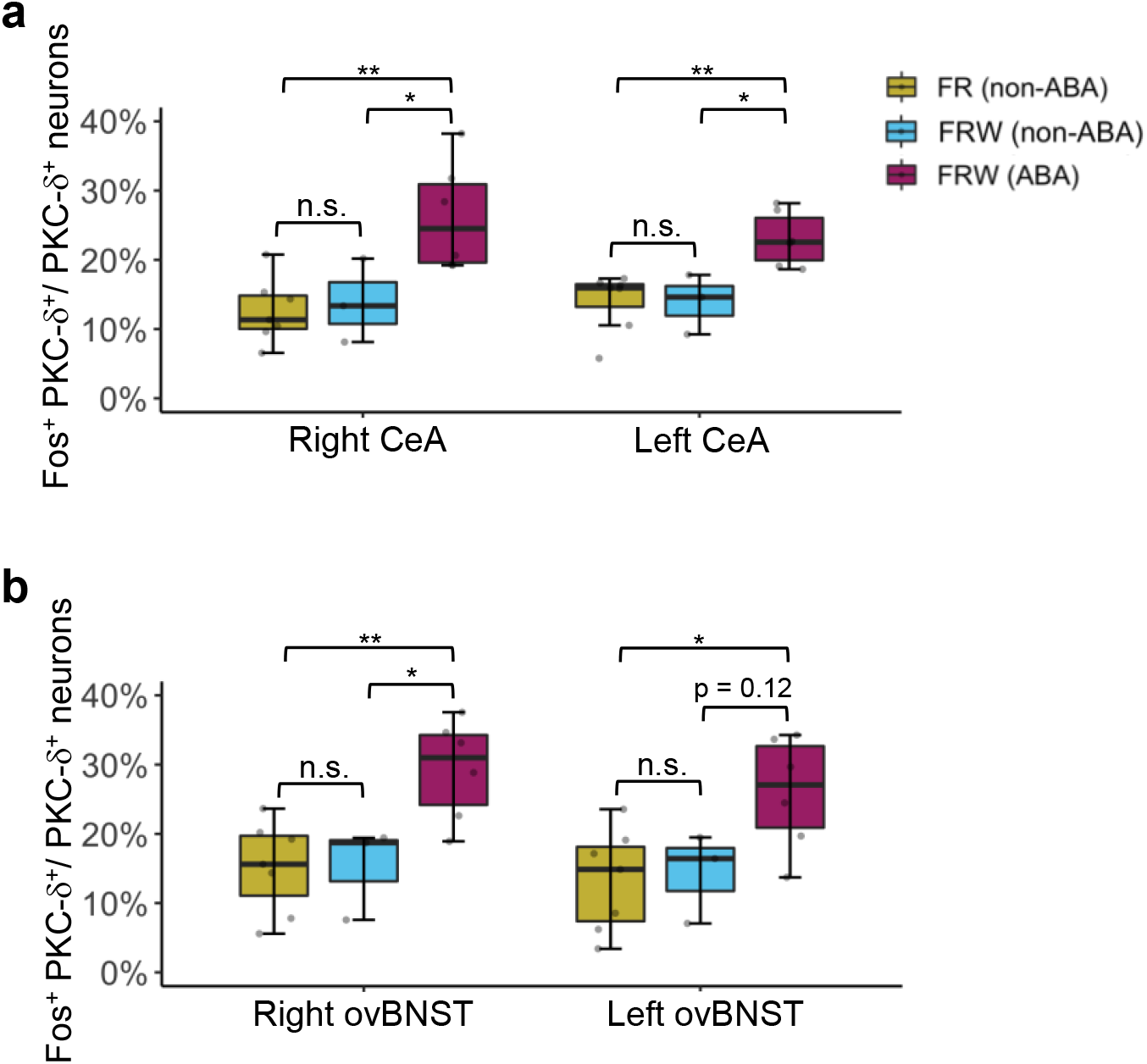
C-Fos expression in EAc^PKC-δ^ neurons is significantly increased bilaterally. **a-b,** Quantification of Fos-like immunoreactivity in both right and left sides of the CeA (**a**) and ovBNST (**b**) brain regions of WT mice in the food restricted only condition (FR Non-ABA; n = 7), ABA resistant mice with food restriction and wheel (FRW Non-ABA; n = 3), and ABA susceptible mice with food restriction and wheel (FRW ABA; n = 6). One-way ANOVA with Tukey HSD post-hoc. All samples are females. * *p* < 0.05, ** *p* < 0.01, *** *p* < 0.001.

**Extended Data Fig. 5.**
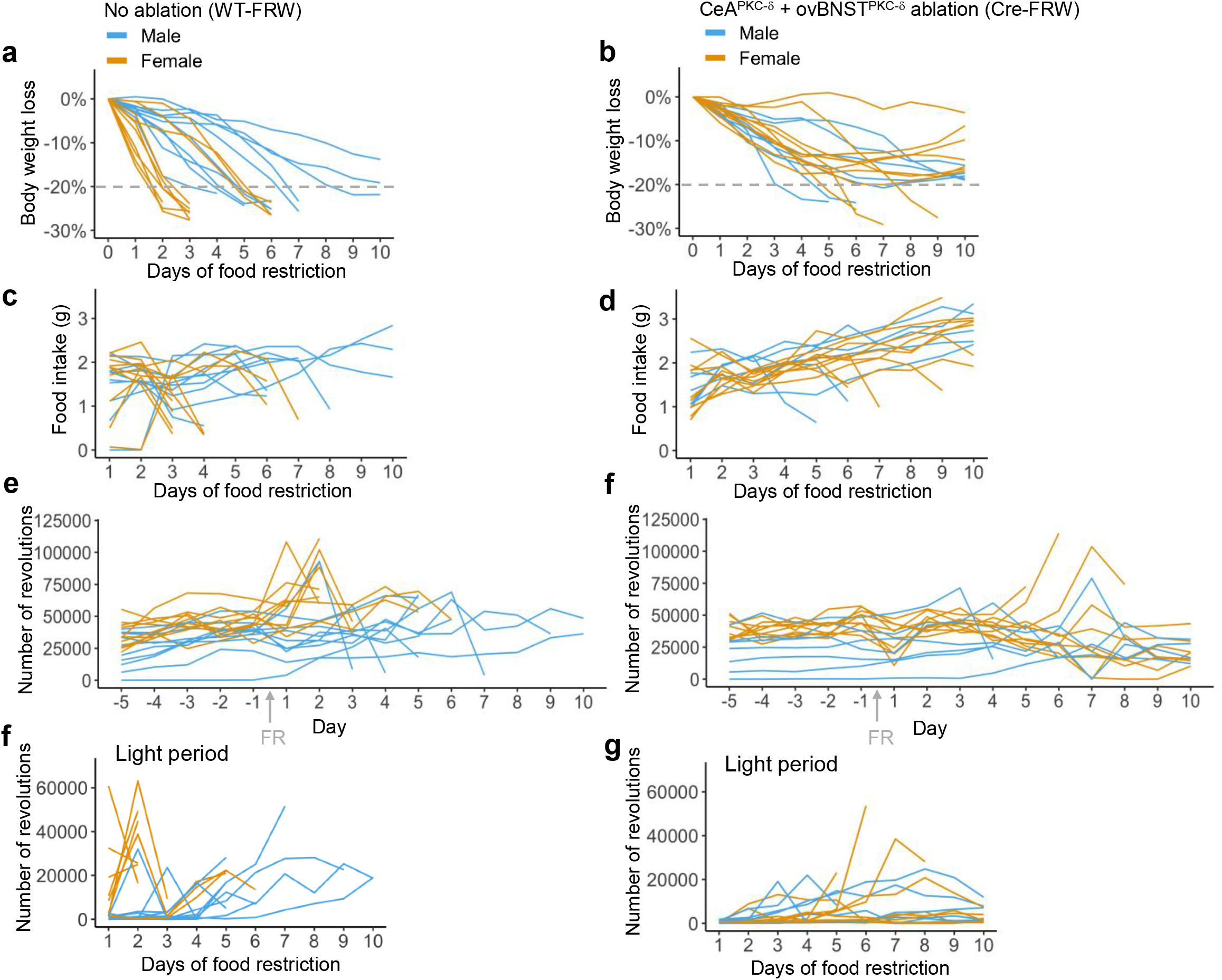
Male and female trends in the ABA paradigm are contrasting for WT-FRW mice, but similar for Cre-FRW mice with dual EAc^PKC-δ^ neuron ablation. **a-d,** Individual sample data across days of food restriction, comparing males and females, for body weight loss (**a-b**), food intake (**c-d**). **e-g,** Total number of wheel revolutions for each mouse 5 days preceding food restriction into the 10 days of food restriction (**e-f**), and during the light period (4 am - 4 pm) on days of food restriction (**f-g**). Left columns: no ablation WT mice. Right columns, dual ablation Cre mice. Arrow in (**e, f**) indicates day in which food restriction was enforced. WT: male n = 10, female n = 10, Cre: male n = 7 and female n = 10.

**Extended Data Fig. 6.**
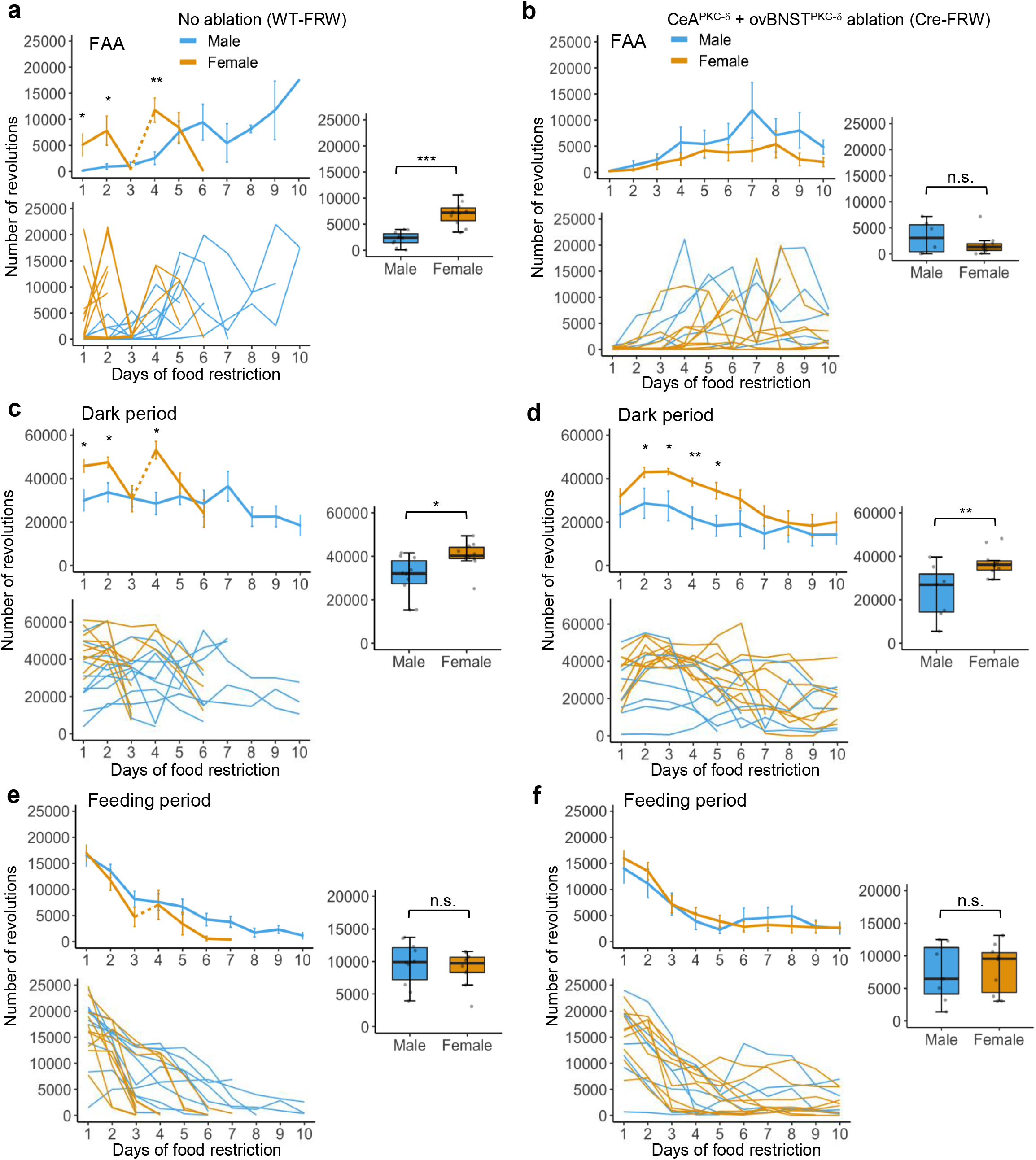
Sexual divergence in food anticipatory activity (FAA) during ABA is diminished with dual EAc^PKC-δ^ neuron ablation. **a-f,** Population mean data (top left), individual samples (bottom left), and average across days 1-5 of food restriction (right) for total wheel activity during food anticipatory activity (FAA, 4 hours preceding food) (**a-b**), dark period (4 pm - 4 am) (**c-d**), and feeding period (**e-f**). Sudden change in the line plots for female WT mice (i.e., days 3-4) is due to the removal of half the initial number of mice from the experiment in one day; indicated by dashed section in line. WT: male n = 10, female n = 10, Cre: male n = 7 and female n = 10. Unpaired *t*-tests. * *p* < 0.05, ** *p*< 0.01, *** *p* < 0.001.

## REFERENCES

1 Arcelus, J., Mitchell, A. J., Wales, J. & Nielsen, S. Mortality rates in patients with anorexia nervosa and other eating disorders. A meta-analysis of 36 studies. Arch Gen Psychiatry 68, 724–731, doi:10.1001/archgenpsychiatry.2011.74 (2011).

2 Jagielska, G. & Kacperska, I. Outcome, comorbidity and prognosis in anorexia nervosa. Psychiatr Pol 51, 205–218, doi:10.12740/PP/64580 (2017).

3 American Psychiatric Association. Desk reference to the diagnostic criteria from DSM-5. (American Psychiatric Publishing, 2013).

4 Guarda, A. S., Schreyer, C. C., Boersma, G. J., Tamashiro, K. L. & Moran, T. H. Anorexia nervosa as a motivated behavior: Relevance of anxiety, stress, fear and learning. Physiol Behav 152, 466–472, doi:10.1016/j.physbeh.2015.04.007 (2015).

5 Hebebrand, J. et al. Hyperactivity in patients with anorexia nervosa and in semistarved rats: evidence for a pivotal role of hypoleptinemia. Physiol Behav 79, 25–37, doi: 10.1016/s0031-9384(03)00102-1 (2003).

6 Kron, L., Katz, J. L., Gorzynski, G. & Weiner, H. Hyperactivity in anorexia nervosa: a fundamental clinical feature. Compr Psychiatry 19, 433–440, doi:10.1016/0010-440x(78)90072-x (1978).

7 Mattar, L., Thiebaud, M. R., Huas, C., Cebula, C. & Godart, N. Depression, anxiety and obsessive-compulsive symptoms in relation to nutritional status and outcome in severe anorexia nervosa. Psychiatry Res 200, 513–517, doi:10.1016/j.psychres.2012.04.032 (2012).

8 Zipfel, S., Giel, K. E., Bulik, C. M., Hay, P. & Schmidt, U. Anorexia nervosa: aetiology, assessment, and treatment. Lancet Psychiatry 2, 1099–1111, doi:10.1016/S2215-0366(15)00356-9 (2015).

9 O’Hara, C. B., Campbell, I. C. & Schmidt, U. A reward-centred model of anorexia nervosa: a focussed narrative review of the neurological and psychophysiological literature. Neurosci Biobehav Rev 52, 131–152, doi:10.1016/j.neubiorev.2015.02.012 (2015).

10 Cai, H., Haubensak, W., Anthony, T. E. & Anderson, D. J. Central amygdala PKC-delta(+) neurons mediate the influence of multiple anorexigenic signals. Nat Neurosci 17, 1240–1248, doi:10.1038/nn.3767 (2014).

11 Wang, Y. et al. A bed nucleus of stria terminalis microcircuit regulating inflammation-associated modulation of feeding. Nat Commun 10, 2769, doi:10.1038/s41467-019-10715-x (2019).

12 Routtenberg, A. & Kuznesof, A. W. Self-starvation of rats living in activity wheels on a restricted feeding schedule. J Comp Physiol Psychol 64, 414–421 (1967).

13 Francois, M. & Zeltser, L. M. Rethinking the Approach to Preclinical Models of Anorexia Nervosa. Curr Psychiatry Rep 24, 71–76, doi:10.1007/s11920-022-01319-2 (2022).

14 Zhang, J. & Dulawa, S. C. The Utility of Animal Models for Studying the Metabo-Psychiatric Origins of Anorexia Nervosa. Front Psychiatry 12, 711181, doi:10.3389/fpsyt.2021.711181 (2021).

15 Gutierrez, E. A rat in the labyrinth of anorexia nervosa: contributions of the activity-based anorexia rodent model to the understanding of anorexia nervosa. Int J Eat Disord 46, 289–301, doi:10.1002/eat.22095 (2013).

16 Kurnik-Łucka, M., Skowron, K., Gil, K. In Search for Perfection: An Activity-Based Rodent Model of Anorexia. Animal Models of Eating Disorders. Second edn, Vol. 161 363–377 (Humana, 2021).

17 Klenotich, S. J. & Dulawa, S. C. The activity-based anorexia mouse model. Methods Mol Biol 829, 377–393, doi:10.1007/978-1-61779-458-2_25 (2012).

18 Schalla, M. A. & Stengel, A. Activity Based Anorexia as an Animal Model for Anorexia Nervosa-A Systematic Review. Front Nutr 6, 69, doi:10.3389/fnut.2019.00069 (2019).

19 Culbert, K. M., Sisk, C. L. & Klump, K. L. A Narrative Review of Sex Differences in Eating Disorders: Is There a Biological Basis? Clin Ther 43, 95–111, doi: 10.1016/j.clinthera.2020.12.003 (2021).

20 Welch, A. C., Katzka, W. R. & Dulawa, S. C. Assessing Activity-based Anorexia in Mice. J Vis Exp, doi:10.3791/57395 (2018).

21 Beneke, W. M., Schulte, S. E. & vander Tuig, J. G. An analysis of excessive running in the development of activity anorexia. Physiol Behav 58, 451–457, doi: 10.1016/0031-9384(95)00083-u (1995).

22 Beeler, J. A. & Burghardt, N. S. Activity-based Anorexia for Modeling Vulnerability and Resilience in Mice. Bio Protoc 11, e4009, doi:10.21769/BioProtoc.4009 (2021).

23 Beeler, J. A. et al. Vulnerable and Resilient Phenotypes in a Mouse Model of Anorexia Nervosa. Biol Psychiatry 90, 829–842, doi:10.1016/j.biopsych.2020.06.030 (2021).

24 Pierce, W. D. et al. Activity anorexia: An interplay between basic and applied behavior analysis. Behav Anal 17, 7–23, doi:10.1007/BF03392649 (1994).

25 Chowdhury, T. G., Chen, Y. W. & Aoki, C. Using the Activity-based Anorexia Rodent Model to Study the Neurobiological Basis of Anorexia Nervosa. J Vis Exp, e52927, doi: 10.3791/52927 (2015).

26 Mistlberger, R. E. Circadian food-anticipatory activity: formal models and physiological mechanisms. Neurosci Biobehav Rev 18, 171–195, doi:10.1016/0149-7634(94)90023-x (1994).

27 Verhagen, L. A., Luijendijk, M. C., Hillebrand, J. J. & Adan, R. A. Dopamine antagonism inhibits anorectic behavior in an animal model for anorexia nervosa. Eur Neuropsychopharmacol 19, 153–160, doi:10.1016/j.euroneuro.2008.09.005 (2009).

28 Achamrah, N. et al. Sex differences in response to activity-based anorexia model in C57Bl/6 mice. Physiol Behav 170, 1–5, doi:10.1016/j.physbeh.2016.12.014 (2017).

29 Bartling, B. et al. Sex-related differences in the wheel-running activity of mice decline with increasing age. Exp Gerontol 87, 139–147, doi:10.1016/j.exger.2016.04.011 (2017).

30 Manzanares, G., Brito-da-Silva, G. & Gandra, P. G. Voluntary wheel running: patterns and physiological effects in mice. Braz J Med Biol Res 52, e7830, doi:10.1590/1414-431X20187830 (2018).

31 Bulik, C. M. et al. Genetics and neurobiology of eating disorders. Nat Neurosci 25, 543–554, doi:10.1038/s41593-022-01071-z (2022).

32 Kaye, W. H., Wierenga, C. E., Bailer, U. F., Simmons, A. N. & Bischoff-Grethe, A. Nothing tastes as good as skinny feels: the neurobiology of anorexia nervosa. Trends Neurosci 36, 110–120, doi:10.1016/j.tins.2013.01.003 (2013).

33 Ross, R. A., Mandelblat-Cerf, Y. & Verstegen, A. M. Interacting Neural Processes of Feeding, Hyperactivity, Stress, Reward, and the Utility of the Activity-Based Anorexia Model of Anorexia Nervosa. Harv Rev Psychiatry 24, 416–436, doi:10.1097/HRP.0000000000000111 (2016).

34 Beeler, J. A. & Burghardt, N. S. Commentary on Vulnerability and Resilience to Activity-Based Anorexia and the Role of Dopamine. J Exp Neurol 2, 21–28 (2021).

35 Klenotich, S. J., Ho, E. V., McMurray, M. S., Server, C. H. & Dulawa, S. C. Dopamine D2/3 receptor antagonism reduces activity-based anorexia. Transl Psychiatry 5, e613, doi: 10.1038/tp.2015.109 (2015).

36 Cai, X. et al. A D2 to D1 shift in dopaminergic inputs to midbrain 5-HT neurons causes anorexia in mice. Nat Neurosci 25, 646–658, doi:10.1038/s41593-022-01062-0 (2022).

37 Welch, A. C. et al. Dopamine D2 receptor overexpression in the nucleus accumbens core induces robust weight loss during scheduled fasting selectively in female mice. Mol Psychiatry 26, 3765–3777, doi:10.1038/s41380-019-0633-8 (2021).

38 Foldi, C. J., Milton, L. K. & Oldfield, B. J. The Role of Mesolimbic Reward Neurocircuitry in Prevention and Rescue of the Activity-Based Anorexia (ABA) Phenotype in Rats. Neuropsychopharmacology 42, 2292–2300, doi:10.1038/npp.2017.63 (2017).

39 Milton, L. K. et al. Suppression of Corticostriatal Circuit Activity Improves Cognitive Flexibility and Prevents Body Weight Loss in Activity-Based Anorexia in Rats. Biol Psychiatry 90, 819–828, doi:10.1016/j.biopsych.2020.06.022 (2021).

40 Sternson, S. M. & Eiselt, A. K. Three Pillars for the Neural Control of Appetite. Annu Rev Physiol 79, 401–423, doi:10.1146/annurev-physiol-021115-104948 (2017).

41 Andermann, M. L. & Lowell, B. B. Toward a Wiring Diagram Understanding of Appetite Control. Neuron 95, 757–778, doi:10.1016/j.neuron.2017.06.014 (2017).

42 Watts, A. G., Kanoski, S. E., Sanchez-Watts, G. & Langhans, W. The physiological control of eating: signals, neurons, and networks. Physiol Rev 102, 689–813, doi:10.1152/physrev.00028.2020 (2022).

43 Miletta, M. C. et al. AgRP neurons control compulsive exercise and survival in an activity-based anorexia model. Nat Metab 2, 1204–1211, doi:10.1038/s42255-020-00300-8 (2020).

44 Murray, S. B. et al. Fear as a translational mechanism in the psychopathology of anorexia nervosa. Neurosci Biobehav Rev 95, 383–395, doi:10.1016/j.neubiorev.2018.10.013 (2018).

45 Hardaway, J. A., Crowley, N. A., Bulik, C. M. & Kash, T. L. Integrated circuits and molecular components for stress and feeding: implications for eating disorders. Genes, brain, and behavior 14, 85–97, doi:10.1111/gbb.12185 (2015).

46 Petrovich, G. D., Ross, C. A., Mody, P., Holland, P. C. & Gallagher, M. Central, but not basolateral, amygdala is critical for control of feeding by aversive learned cues. J Neurosci 29, 15205–15212, doi:29/48/15205 [pii] 10.1523/JNEUROSCI.3656-09.2009 (2009).

47 Jennings, J. H., Rizzi, G., Stamatakis, A. M., Ung, R. L. & Stuber, G. D. The inhibitory circuit architecture of the lateral hypothalamus orchestrates feeding. Science 341, 1517–1521, doi:10.1126/science.1241812 (2013).

48 Douglass, A. M. et al. Central amygdala circuits modulate food consumption through a positive-valence mechanism. Nat Neurosci 20, 1384–1394, doi:10.1038/nn.4623 (2017).

49 Kim, J., Zhang, X., Muralidhar, S., LeBlanc, S. A. & Tonegawa, S. Basolateral to Central Amygdala Neural Circuits for Appetitive Behaviors. Neuron 93, 1464–1479 e1465, doi:10.1016/j.neuron.2017.02.034 (2017).

50 Hardaway, J. A. et al. Central Amygdala Prepronociceptin-Expressing Neurons Mediate Palatable Food Consumption and Reward. Neuron 102, 1037–1052 e1037, doi:10.1016/j.neuron.2019.03.037 (2019).

51 Ip, C. K. et al. Amygdala NPY Circuits Promote the Development of Accelerated Obesity under Chronic Stress Conditions. Cell Metab 30, 111–128 e116, doi:10.1016/j.cmet.2019.04.001 (2019).

52 Griessner, J. et al. Central amygdala circuit dynamics underlying the benzodiazepine anxiolytic effect. Mol Psychiatry 26, 534–544, doi:10.1038/s41380-018-0310-3 (2021).

53 Haubensak, W. et al. Genetic dissection of an amygdala microcircuit that gates conditioned fear. Nature 468, 270–276 (2010).

54 Mistlberger, R. E. Food as circadian time cue for appetitive behavior. F1000Res 9, doi:10.12688/f1000research.20829.1 (2020).

55 Amir, S. & Stewart, J. Behavioral and hormonal regulation of expression of the clock protein, PER2, in the central extended amygdala. Prog Neuropsychopharmacol Biol Psychiatry 33, 1321–1328, doi:10.1016/j.pnpbp.2009.04.003 (2009).

56 Perrin, J. S., Segall, L. A., Harbour, V. L., Woodside, B. & Amir, S. The expression of the clock protein PER2 in the limbic forebrain is modulated by the estrous cycle. Proc Natl Acad Sci U S A 103, 5591–5596, doi:10.1073/pnas.0601310103 (2006).

57 Ye, J. & Veinante, P. Cell-type specific parallel circuits in the bed nucleus of the stria terminalis and the central nucleus of the amygdala of the mouse. Brain Struct Funct 224, 1067–1095, doi:10.1007/s00429-018-01825-1 (2019).

58 Davis, M., Walker, D. L., Miles, L. & Grillon, C. Phasic vs sustained fear in rats and humans: role of the extended amygdala in fear vs anxiety. Neuropsychopharmacology 35, 105–135, doi:10.1038/npp.2009.109 (2010).

59 Kim, S. Y. et al. Diverging neural pathways assemble a behavioural state from separable features in anxiety. Nature 496, 219–223, doi:10.1038/nature12018 (2013).

60 Cassell, M. D., Freedman, L. J. & Shi, C. The intrinsic organization of the central extended amygdala. Ann N Y Acad Sci 877, 217–241 (1999).

61 Yang, C. F. et al. Sexually dimorphic neurons in the ventromedial hypothalamus govern mating in both sexes and aggression in males. Cell 153, 896–909, doi:10.1016/j.cell.2013.04.017 (2013).

62 Franklin, K. B. J. & Paxinos, G. The Mouse Brain in Stereotaxic Coordinates. Third edn, (Academic Press is an imprint of Elsevier, 2007).

